# The temporal copulatory patterns of female rat sexual behavior

**DOI:** 10.1101/2024.10.07.616964

**Authors:** John C. Oyem, Roy Heijkoop, Eelke MS Snoeren

## Abstract

Female sexual behavior is a naturally rewarding activity that plays an important role in reproduction and species survival. For female rats, regulating the timing of sexual interactions is essential for optimizing mating satisfaction and enhancing the physiological conditions needed for successful fertilization. So far, traditional research on female sexual behavior has relied on a limited set of behavioral parameters, which has certain shortcomings. To address this, our study aimed to develop a more detailed behavioral framework for assessing temporal copulatory patterns in female rats. We compared fully receptive females and less-receptive females, while also investigating the effects of (R)-(+)-8-OH-DPAT, a 5-HT1A receptor agonist known for its inhibitory impact on female sexual behavior. Additionally, we examined how sexual experience and pacing conditions influence these copulatory patterns. Our results revealed that female rats engage in structured patterns of sexual bouts and time-outs, with higher receptivity leading to more sexual bouts and shorter time-outs. This suggests that sexual bouts can be viewed as an indicator of copulatory intensity, while time-outs reflect motivation to continue mating. Sexual experience did not enhance sexual performance but did result in females receiving more copulatory events from males. Lastly, we found that the conditions under which mating occurs (paced vs. non-paced) may not significantly impact copulatory behavior in fully-receptive females but could be more relevant for less-receptive females. Despite this, paced mating conditions remain preferable for studying female sexual behavior.

## 1 Introduction

Female sexual behavior is a natural rewarding behavior that plays an essential role in reproduction and the survival of the species. From observations in the field and more controlled experiments in semi-natural environments, we learned that rats often mate in groups, where one or multiple males and females interact simultaneously (Robitaille and Bovet 1976; Calhoun 1963; Chu and Ågmo 2014; Hegstad et al. 2020). Repetitive patterns of paracopulatory behaviors (such as darts and hops) and reflexive lordosis responses to mounts, intromissions, or ejaculations represent the sexual behaviors of the female rat. In these natural settings, the presence of multiple receptive females and the availability of space for females to move away from sexually active males give them the ability to control, or “pace,” their mating encounters. In the lab, however, the use of small copulation boxes limits females to express this kind of pacing activities, which is why this is referred to as non-paced mating. To overcome this limitation, a divider with small holes, that limit passage to the smaller sized female animals, can be placed inside the copulation box, allowing the female to withdraw to a separate compartment. In this paced mating set-up, the female regains control over the timing and number of sexual encounters with the male rat. Studies in the lab have shown that by managing the intervals between these interactions, female rats can optimize the pleasure derived from mating (Erskine and Hanrahan 1997; Martinez and Paredes 2001) and the physiological conditions necessary for successful fertilization (Coopersmith and Erskine 1994; Erskine 1989). Understanding these dynamics in behavior can provide important knowledge into how female-driven sexual behavior influences reproductive success and ensures the continuation of the species. Moreover, analyzing these complex patterns of behavior become essential when studying the neurobiological mechanisms underlying mating (Krakauer et al. 2017).

Traditionally, a limited amount of parameters has been used to analyze and interpret female sexual behavior. While the number of paracopulatory behaviors may reflect the level of sexual motivation, sexual receptivity is measured with a lordosis quotient and score (reviewed in (Heijkoop, Huijgens, and Snoeren 2018; Ventura-Aquino and Paredes 2023)). In a paced-mating setting, the withdrawal behavior can be analyzed by calculating the percentages of exits after mounts, intromissions and ejaculations and the time it takes for the female to resume copulation (also called contact return latency). These pacing parameters, as they are also called, are dependent on the stimulation intensity, with the largest percentages and longest return latencies after ejaculation, followed by intromissions and mounts (Krieger, Orr, and Perper 1976; Meerts et al. 2014; Brandling-Bennett, Blasberg, and Clark 1999; Zipse, Brandling-Bennett, and Clark 2000; Erskine 1985; Erskine 1989). While the percentages of exits may indicate the female rats’ ability to discriminate sensory stimulation, contact return latencies (CRL) reflect her motivation to continue copulation (reviewed in (Heijkoop, Huijgens, and Snoeren 2018)).

While percentages of exits and CRL offer interesting insights into the pacing behavior of female rats, they also have limitations. For instance, CRL can only be measured following an exit, which excludes latencies where the female immediately resumes copulation. Furthermore, there is variability in how different laboratories define an exit. Some labs set a specific time limit within which the female must have moved to the other compartment to be considered an exit, while others define an exit without considering a withdrawal time. The question therefore remains what a CRL actually represents. Finally, as the females do not have control over the type of stimulation they receive, their pacing behavior is highly dependent on the behavior of the male. It has been shown, for example, that CRL after ejaculations, sometimes referred to as female post-ejaculatory intervals (PEI), are longer when the same male remains in the mating chamber than when a different male was introduced (Corlett et al. 2022).

Given the limitations of current parameters used to study female sexual behavior, an improved behavioral assessment tool could be beneficial. Additionally, a more detailed analysis of how females control the timing of sexual encounters beyond a simple withdrawal to another compartment, would do justice to the complexity of this phenomenon. An example can be taken from male rats, for which we recently employed an idea by (Sachs and Barfield 1970), who demonstrated that male rat copulation is temporally organized in mount bouts. These bouts are defined as sequences “of mounts (one or more), with or without intromission, uninterrupted by any behavior (other than genital autogrooming) that is not oriented towards the female.” The breaks between these mount bouts are defined as “time-outs” (Huijgens et al. 2021). Using these parameters in an extended behavioral analyses, we have been able to obtain valuable insights into the neural regulation of male rat sexual behavior, and specific roles for different brain regions have been revealed with the medial amygdala being involved in regulating copulatory sensitivity and ejaculatory threshold (Huijgens, Heijkoop, and Snoeren 2021) and the bed nucleus of stria terminalis in regulating motivation to continue copulation (Huijgens et al. 2024). A similar assessment tool has not yet been developed for female rats, but could potentially provide valuable information about female sexual behavior as well.

Therefore, in this study we aimed to develop a similar behavioral assessment tool for female rats by investigating the temporal copulatory patterns in more detail. We took advantage of the fact that the sexual activity of females is highly dependent on hormonal status (Snoeren, Chan, et al. 2011; Brandling-Bennett, Blasberg, and Clark 1999; Hliňák 1986; Dominguez-Ordonez et al. 2015; Edwards and Pfeifle 1983), which allowed us to compare the temporal patterns of normally receptive females with low receptive females. By using a within-subject design, we were able to follow the progress of each rat over multiple copulation tests and study the effects of sexual experience on their temporal copulatory patterns. Additionally, as an extra proof of concept, we tested the rats once more after treatment with (R)-(+)-8-OH-DPAT, a 5-HT1A receptor agonist. This pharmaceutical compound is known to inhibit paracopulatory behavior (Uphouse and Wolf 2004; Kishitake and Yamanouchi 2003; Mendelson and Gorzalka 1986; Snoeren et al. 2010; Snoeren, Refsgaard, et al. 2011; Snoeren et al. 2014), and could therefore provide information about how inhibition of female sexual behavior would be reflected in the temporal copulatory patterns, and thus help us assess the usefulness and interpretation of bout-based assessments. Finally, we added an experiment in which we investigated whether a paced mating condition would still be important in assessing female sexual behavior when using this new assessment tool for studying female sexual behavior.

## 2 Materials and Methods

### 2.1 Animals

The experiment involved twenty-four sexually naïve adult female and twelve male Wistar rats, each weighing approximately 230g, purchased from Janvier labs, France. Rats were housed in same-sex pairs in Macrolon IV ® homecages in a room with a reversed 12 h light/dark cycle (lights on between 23:00 and 11:00), controlled temperature (21 ± 1 °C), and humidity (55 ± 10%). They were provided standard rodent food pellets (low phytoestrogen maintenance diet, #V1554, Ssniff, Germany) and tap water *ad libitum*.

After one week of acclimatization to the animal facility, female rats were ovariectomized under isoflurane anesthesia following the methods described by (Ågmo 1997). A medial dorsal incision of about 1 cm long was made in the skin. Using blunt dissection, abdominal musculature was exposed just enough to allow a bilateral incision of about 0.5 cm in the dorsolateral abdominal musculature. The ovaries on each side were located and extirpated, while the fallopian tubes were ligated and gently placed back in the posterior abdomen. The incised muscle layer was closed with absorbable sutures (Vicryl 4-0, Ethicon), and the skin was closed using wound clips. The male rats, on the other hand, were sexually trained by allowing them to copulate for 30 minutes on three occasions with receptive females who are not further used or mentioned in this study. The sexually trained rats were then used as stimulus males in this study.

All animal care and experimental procedures employed in this study were conducted in agreement with the European Union Council Directive 2010/63/EU and in accordance with the Norwegian Food Safety Authority.

### 2.2 Experimental groups (models)

The ovariectomized female rats were randomly assigned to a hormonally sub-primed (n=12) and hormonally fully-primed group (n=12). The hormonally sub-primed females were administered 5 μg of Estradiol benzoate (EB, Sigma-Aldrich, product Nr: E8875) alone, while hormonally fully-primed females were administered 5 μg of EB and 500 μg of progesterone (P, Sigma-Aldrich, product Nr: P-0130). Both EB and P were dissolved in peanut oil solution at 5 mg/mL. EB and P were administered subcutaneously at different time intervals: EB was given 36 hours before the start of each copulation session, while P was administered 4 hours prior to each copulation session.

The administration of EB alone induces lower receptivity than the administration of the combination of EB and P, which induces full receptivity in female rats (Snoeren, Bovens, et al. 2011). According to a previous study (Snoeren, Bovens, et al. 2011), sub-primed females are intended to represent females with low receptivity, potentially modeling female sexual dysfunction, while the fully-primed group is used to model normally receptive females.

### 2.3 Apparatus

We used the paced and non-paced mating apparatus in this experiment. The paced mating apparatus consists of a rectangular steel box measuring 40 x 60 x 40 cm with a Plexiglas front and an arena covered with wood chips. The interior space is divided into two compartments by a transparent compartment divider. The divider has three 4 cm diameter holes at the bottom, which serve as an escape route for the females during copulation. As a result, one compartment (40 x 45 x 40 cm) is available for both the male and female, while a smaller compartment (40 x 15 x 40 cm) can only be accessed by the female. For the non-paced mating, we removed the transparent compartment divider in the paced mating apparatus, transforming the apparatus into one compartment (40 x 60 x 40 cm) where rats can copulate without an escape route for the females.

### 2.4 Behavioral testing

All copulation sessions started at noon and were conducted in a room with a 5 lux dim light. The experimenter was blinded to the experimental group of the subject animals. The female rats were placed in the female compartment and were allowed to habituate in the copulation apparatus for five minutes. Next, a stimulus male rat was placed in the male compartment, and they were allowed to copulate for 30 minutes. If during that time the stimulus male achieved an ejaculation, he was replaced with another stimulus male (the two males were circulated within the same copulation session). This was done to enable us to study the female rats’ copulatory pattern independent of the male’s post-ejaculatory interval. The entire copulation test was videotaped, and behavioral assessment was carried out by annotating all female sexual behaviors with the use of observer XT version 17 software (Noldus, Wageningen, the Netherlands).

### 2.5 Experimental setup

#### 2.5.1. Experiment part 1

The goal of the first part of the experiment was to determine the temporal copulatory patterns of female rats and the effect of gaining sexual experience on these female temporal copulatory patterns. Two groups of sexually naïve female rats, sub-primed and fully-primed, were exposed every 5^th^ day to a copulation test in the paced mating setup for 6 times (Cop 1 to Cop 6) (Fig 1A). With this setup, we explored the temporal copulatory patterns of the female rats and the effects of sexual experience by comparing the data from naïve (Cop 1) versus sexually experienced (Cop 6) in sub-and fully-primed rats. To obtain the progress of gaining sexual experience, also Cop 2 and Cop 4 were analyzed.

#### 2.5.2. Experiment part 2

In the second part of the experiment, we tested the hypothesis that (R)-(+)-8-OH-DPAT (Sigma-Aldrich, product Nr: H140-5MG), a 5-HT1A receptor agonist known to inhibit female sexual behavior (Snoeren et al. 2010), could disrupt the temporal copulatory patterns in females. For this purpose, we used all female rats from the 1^st^ part of the experiment in part 2 (Fig. 1A). The copulation test was performed twice in a within-subject Latin square design (with a 1-week interval, which served as a wash-out period). We administered 0.1 mg/kg of (R)-(+)-8-OH-DPAT or vehicle subcutaneously 10 minutes before the copulation test in a paced mating setup.

#### 2.5.3. Experiment part 3

Finally, the same female rats were tested in a non-paced mating setup for 30 minutes. The aim of the last part of the experiment was to compare the temporal copulatory pattern of female rats in paced mating versus non-paced mating. With this, we can infer whether the female’s temporal copulatory pattern is dependent on the setup in which she can withdraw from the male and thus actively pace her sexual encounters versus the setup in which she is not able to withdraw from the male during copulation. The same animals from experimental parts 1 and 2 were also used in part 3. Two non-paced mating tests were performed twice to familiarize the rats with the new concept (Fig 1A). Only the last non-paced mating test was used for further analysis.

### 2.6 Behavioral analysis

We manually scored the following female behaviors: paracopulatory behaviors (hops and darts), female receptive behavior (lordosis behavior on a 4-point scale (Hardy and Debold 1971)), sniffing (sniffing other body parts of the male), anogenital sniffing (sniffing the anogenital region of the male), genital self-grooming, other behaviors such as female running, head towards the male (head oriented in the direction of the male), and head away from the male (head oriented away from the male). The male behaviors that were annotated included mounts, intromissions, and ejaculations. In addition, we scored the occasions in which the female crossed over between compartments, so that we could calculate pacing behavior measures such as: the percentage of exits following copulations (mounts, intromissions, and ejaculations), contact return latencies following exits, and the total time spent in each compartment.

Furthermore, using a similar approach as previously done for male rats (Sachs and Barfield 1970; Huijgens et al. 2021), we divided the female rats’ temporal copulatory patterns into sexual bouts and time-outs. A female sexual bout was defined as a series of behaviors that begins and ends with either a paracopulatory behavior or lordosis response. All subsequent behaviors oriented towards the male (such as sniffing, anogenital snigging, genital self-grooming, head towards the male) were part of the same bout, until a behavior was displayed that was not oriented towards the male (such as head away from the male, or rejection behaviors). Then the former last paracopulatory or lordosis behavior marked the end of a sexual bout, and a time-out was started and lasted until the next paracopulatory behavior or lordosis response (Fig. 2A). A self-written Python script was used to analyze all sexual behaviors and identify the female sexual bouts. These behavioral outcome measures are listed in Suppl. Table 1.

### 2.7 Statistical analysis

Statistical analyses were conducted using SPSS software (version 29, IBM, Armonk, USA) with a level for statistically significant difference set at p < 0.05. A Shapiro-Wilk test confirmed that the data was not normally distributed, and therefore a linear mixed model was used that comprised of the factors (hormonal) treatment and experience for the sexual experience study (part 1), treatment and drug for the (R)-(+)-8-OH-DPAT study (part 2), and treatment and pacing for the paced and non-paced mating study (part 3). In the case of a significant interaction effect, a Bonferroni-corrected post hoc test was conducted. When a correlation test was used to analyze the relationship between the duration of sexual bouts and time-outs and other parameters, the Spearman correlation test was conducted.

Finally, in order to investigate what determines the duration of sexual bouts and time-outs, data points were z-scored within each rat using the following calculation: z-score = ((data point) - (mean of the data points for the rat)) / (standard deviation of the data points for the rat). Z-scores were then analyzed by means of the Kruskal-Wallis test, followed by Mann Whitney U post hoc tests.

## 3 Experiment part 1

In the first part of the experiment, we attempted to develop an improved behavioral assessment tool by studying the female rats’ temporal copulatory patterns in detail. In addition, we investigated the effects of sexual experience on these temporal copulatory patterns. By using sub- and fully-prime females, the effect of low versus normal receptivity, respectively, could be assessed.

To make this result section more readable, we moved all statistical outcomes on the hormonal treatment, experience, and/or interaction effects into a separate table (Suppl. Table 2) and report only on the significant relevant results in this paper.

### 3.1 Results traditional parameters

When looking at the more traditional parameters first, our data revealed that sub-primed females spent a similar amount of time in the male compartment as fully-primed females, and that this time decreases after a single sexual experience (Fig. 1B). Additionally, sub-primed females exhibited fewer paracopulatory behaviors compared to fully-primed females across all copulation (Cop) tests (Fig. 1C), including when only paracopulatory behaviors performed in the female compartment were considered (Suppl. Figure S1). Furthermore, sub-primed females demonstrated a longer latency to their first paracopulatory behavior (measured from the male’s introduction) than fully-primed females, particularly in the first copulation (Cop1) test (Suppl. Table 3). Although sexual experience did not alter the overall number of paracopulatory behaviors, it did increase the number of darts displayed in the female compartment for both sub- and fully-primed females. Additionally, sexual experience reduced the latency to the first paracopulatory behavior in sub-primed females, bringing it in line with fully-primed females.

Regarding lordosis behavior, we found that sexually naïve (Cop 1) fully-primed females had a significantly higher lordosis quotient (LQ, calculated as number of lordosis responses divided by number of received copulations times 100%) than sub-primed females (Fig. 1D). This difference, however, disappeared after the females gained sexual experience. The lordosis score (LS, intensity of the lordosis response), on the other hand, seemed to increase in Cop6 compared to Cop1 in fully-primed females, but no differences were found between sub- and fully primed females (Fig. 1E). It should be mentioned, though, that two rats were removed from the data analysis regarding lordosis behavior. This was done because they displayed an extraordinary number of lordoses without receiving any copulations and were therefore considered statistical outliers.

**Figure 1.**
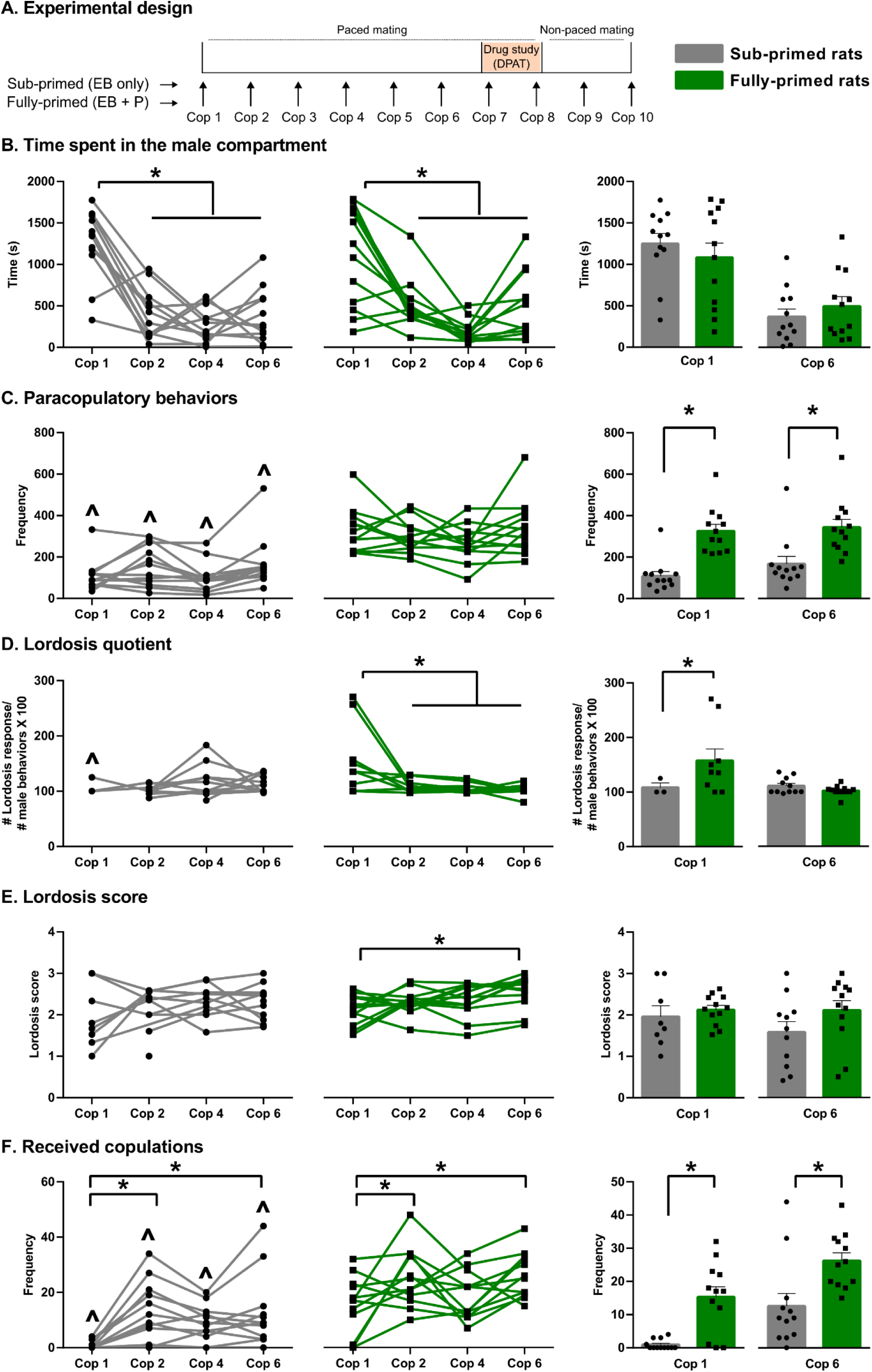
Sexual experience and hormonal status and female rats copulatory parameters. (A) Schematic illustration of the experimental timeline, (B) Time spent in the male compartment (in seconds), (C) Total number of paracopulatory behaviors, (D) Lordosis quotient, (E) Lordosis score, (F) Total number of copulations (mounts, intromission and ejaculations) received by the females. **Panels to the left (B-F):** The data are shown with individual data points, with the lines connecting each rat across different Cop tests; panels **to the right (B-F)**: The data are shown with individual data points, with the bars representing the mean ± SEM, **All figures (B-F):** Data is shown for sub- and fully-primed female Wistar rats. *p < 0.05 significantly different between Cop tests, ^ʌ^p < 0.05 is significantly different between groups (sub-vs fully-primed). Cop = Copulation test, EB = Estradiol benzoate, P = progesterone, DPAT = (R)-(+)-8-OH-DPAT.

When exploring the number of received copulations, we found that fully-primed rats received significantly more copulations compared to sub-primed rats in all Cop tests (Fig. 1F). Both sub- and fully-primed rats showed an increase in received copulations in the 2^nd^ and 6^th^ Cop tests compared to Cop 1, driven primarily by the number of received intromissions, as the number of received mounts did not vary with experience (Suppl. Table 3). Interestingly, although the stimulus male rats were sexually experienced from the start of the experiment, both sub- and fully-primed females received significantly fewer ejaculations in the sexually naïve versus experienced state. This suggests that female behavior may play a role in regulating male behavior. Furthermore, when comparing the latency to 1st ejaculation calculated from the 1st paracopulatory behavior, we observed that fully-primed females reached their 1st ejaculation more quickly than sub-primed rats in all Cop tests (Suppl. Table 3). Additionally, sexual experience reduced the latency to 1st ejaculation in both sub- and fully-primed females, with sexually experienced rats achieving ejaculation faster than their sexually naïve counterparts. Still, no differences in number of received ejaculations were found between sub- and fully-primed females (Suppl. Table 3).

Additionally, we investigated the effects of hormonal status and sexual experience on traditional paced-mating behaviors such as the percentage of exits and contact-return latencies. No differences were found between sub-and fully-primed females with regard to percentages of exits after mounts, intromissions, or ejaculations (Suppl. Table 3), nor was there an effect of sexual experience on the percentage of exits after mounts or ejaculations. The level of sexual experience did increase the percentage of exits after intromissions, but post-hoc analysis revealed that this effect was only found in fully-primed rats. With the generally low percentages of exits, the number of data points to determine the contact-return latencies (CRL) became rather small, and no effect of hormonal status or experience was found on CRL after mounts, intromissions, or ejaculations (Suppl. Table 3)

### 3.2 Results sexual bouts and time-outs

With the original scope of this paper of determining whether females (just like males) copulate in so-called sexual bouts, we next examined the female rat’s copulatory patterns into sexual bouts and time-outs. As a reminder, paracopulatory or lordosis behaviors marked the start of a sexual bout, and all subsequent behaviors oriented towards the male are part of the same bout. As soon as a behavior was displayed that was not oriented towards the male, the former last paracopulatory or lordosis behavior marked the end of a sexual bout and the start of the time-out (Fig. 2A).

We found that fully-primed females copulated with more sexual bouts than sub-primed females in all Cop test (except Cop 2, Fig. 2B), a phenomenon that was seen both with and without counting single, isolated paracopulatory behaviors as a sexual bout. Additionally, when the mean duration of these sexual bouts was calculated, we found that in all Cop tests (except Cop 6), fully-primed rats also spent on average more time in a sexual bout than sub-primed rats (Fig. 2C). While the number of sexual bouts was also found to remain stable from sexually naïve to experienced states, the mean duration of a sexual bout declined upon gaining sexual experience in fully-primed rats. No effect of experience, however, was found in sub-primed rats. Similar differences were found in the mean duration of the time-outs between the sexual bouts. Fully-primed females had shorter time-outs in all Cop tests compared to sub-primed rats (Fig 2D). In addition, the mean duration of time-outs did not change upon gaining sexual experience and remained stable over the course of the copulation tests in sub- and fully-primed rats (except for an unexplainable increase in time-out duration in Cop 4 in sub-primed rats).

Analysis of the sexual bouts in a more detailed manner revealed that sexual bouts of fully-primed females consisted of more paracopulatory behaviors (considering only the sexual bouts that contained two or more paracopulatory or lordosis behaviors) than sub-primed rats in all Cop tests, and more lordosis responses in Cop1 and Cop6 (Suppl. Table 3). While the number of paracopulatory behaviors per sexual bout generally remained stable over the course of the copulation tests, a slight increase in the mean number of lordoses responses per sexual bout was found after obtaining sexual experience in sub-, but not fully-primed rats (Suppl. Table 3). Since lordosis is most often a stereotypical response upon a received copulatory stimulation from the male, it is then not surprising that the mean number of copulation behaviors within these sexual bouts follows a similar pattern as the lordosis responses (Suppl. Table 3).

**Figure 2.**
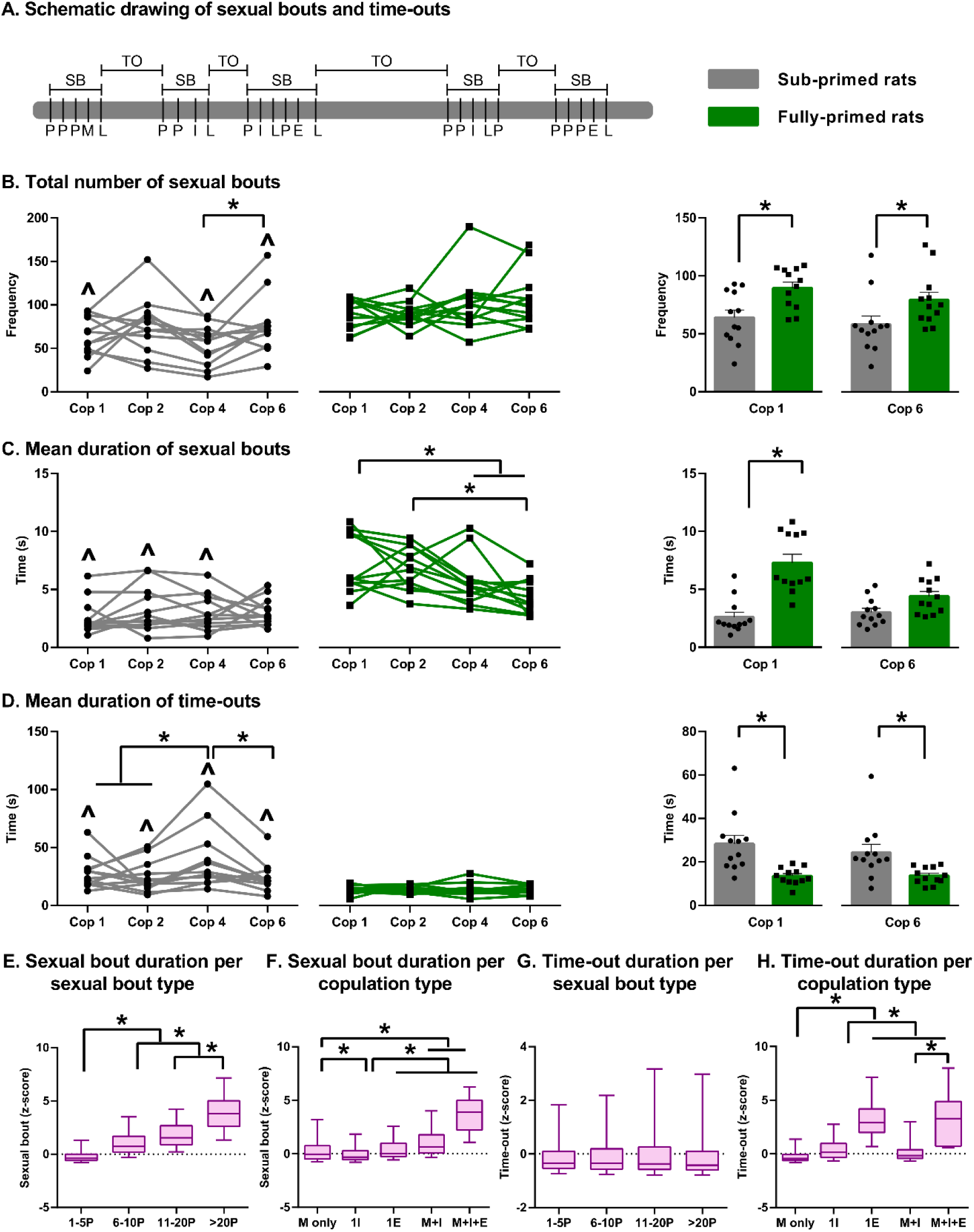
Female temporal copulatory patterns are organized into sexual bouts and time-outs. (A) Schematic representation of female rat temporal copulatory patterns divided into sexual bouts (SB) and time-outs (TO) with paracopulatory behavior (P) and lordosis (L) marking the start and the end of sexual bouts. M = mount, I = intromission, E = ejaculation, (B) Total number of sexual bouts. (C) Mean duration of sexual bouts (in seconds). (D) Mean duration of time-outs (in seconds). (E) Boxplot of z-scores of individual sexual bouts duration divided into groups based on the numbers of paracopulatory behaviors within the bout, (F) Boxplot of z-scores of individual sexual bouts duration divided into groups based on the received copulations within the bout. (G) Boxplot of z-scores of individual time-out duration divided into groups based on the numbers of paracopulatory behaviors within the preceding bout, (H) Boxplot of z-scores of individual time-out duration divided into groups based on the received copulations within the preceding bout. **Panels to the left (B-D):** The data are shown with individual data points, with the lines connecting each rat across different Cop tests; panels **to the right (B-D):** The data are shown with individual data points, with the bars representing the mean ± SEM, **Figures B-D:** Data is shown for sub- and fully-primed female Wistar rats. *p < 0.05 significantly different between tests, **^ʌ^**p < 0.05 is significantly different between groups (sub- vs fully-primed). **Figures E-H:** *p < 0.05 significantly different between sexual bout groups. Cop = Copulation test. P=paracopulatory behavior, M=mount, I=intromission, E=ejaculation.

To understand the newly proposed behavioral analysis of sexual bouts and time-outs better, we investigated what determines the length of these bouts and time-outs. Therefore, we z-scored the durations of each bout and time-out per rat and divided the z-scored bouts and time-outs into groups based on the behaviors that were included in the related sexual bout. As such, to determine the influence of the number of paracopulatory behaviors on the duration of the sexual bouts (without taking into account the copulatory behaviors), the data was divided into four groups: bouts containing 1-5, 6-10, 11-20, and more than 20 paracopulatory behaviors. We found that increasing numbers of paracopulatory behaviors per bout did increase the sexual bouts duration (Fig 2E). Interestingly, when we evaluated the role of different kinds of received copulations during the sexual bouts, we found that the sexual bout durations increased when more intromissions and ejaculations were received, in comparison to only mounts or just 1 intromission or ejaculation (Fig. 2F). It is thus the increased number of copulation within the sexual bout, rather than the type of stimulation that lengthens the sexual bout. Since these copulations often occur with more paracopulatory behaviors, this suggests that the length of a sexual bout is most likely determined by the number of paracopulatory behaviors.

Regarding what determines the length of the time-outs, we found that the duration of time-outs is negatively correlated, but only weakly, with the duration of sexual bouts (data not shown), meaning that longer sexual bouts result in shorter time-outs. Since longer sexual bouts generally contain more paracopulatory behaviors, we next determined the influence of the number of paracopulatory behaviors on the duration of the time-outs (not taking into account the copulatory behaviors). We found that increased numbers of paracopulatory behaviors per bout did not affect the length of the time-outs (Fig 2G). Interestingly, when we evaluated the role of different kinds of received copulations during the sexual bouts, we found that already 1 intromission in the sexual bout lengthens the time-out that follows compared to when the bout consisted of only mounts (Fig 2H). When an intromission was accompanied by mounts or more intromissions, even longer time-outs were seen compared to bouts with only 1 mount, but these time-outs were shorter than time-outs following sexual bouts with 1 intromission. The longest time-outs were found when an ejaculation was received (independent of the co-occurrence of more mounts and/or intromissions within the same sexual bout). Overall, this suggests that while the length of the sexual bout is most likely determined by the number of paracopulatory behaviors, the time-outs are mostly determined by the type of received stimulation and lengthened as soon as an ejaculation was included in the sexual bout.

### 3.3 Discussion Experiment part 1

In summary, our findings support our hypothesis that female rats copulate in temporal copulatory patterns organized into sexual bouts and time-outs. The copulatory patterns are dependent on the hormonal status of the rats, as fully-primed rats copulated with more and longer sexual bouts and shorter time-outs than lower receptive sub-primed females. Sexual experience, on the other hand, did affect the temporal copulatory patterns by shortening the mean duration of the sexual bouts in fully-primed experienced females (to the level of sub-primed females), without changing the total number of sexual bouts and time-out durations. As expected, fully-primed females performed more paracopulatory behaviors and received more male copulations than sub-primed rats. While fully-primed females had a higher LQ than sub-primed rats in Cop1, but not in other copulation tests, no differences were found on LS or pacing behaviors such as exits and CRL after received copulations. Besides a decrease in the lordosis quotient and a small increased the lordosis score in fully-primed rats, sexual experience did not affect other female sexual performance parameters in sub- and fully-primed rats. Sexual experience did only result in receiving more copulations from the second copulation test compared to the 1^st^ in both sub- and fully-primed rats.

In our study, sub-primed rats were hormonally primed with estradiol benzoate (EB) alone, while fully-primed females were primed with estradiol benzoate and progesterone (EB + P). Although EB alone is known to induce hormonal receptivity in female rats, P has been shown to facilitate the effect of EB on e.g. paracopulatory, approach, and lordosis behaviors in female rats (Snoeren, Chan, et al. 2011; Brandling-Bennett, Blasberg, and Clark 1999; Hliňák 1986; Dominguez-Ordonez et al. 2015; Edwards and Pfeifle 1983). Our findings concerning the difference between sub- and fully-primed females on the traditional parameters of paracopulatory behaviors and lordosis quotient and score are therefore in line with these previous studies.

Notably, we did not find any effect of hormonal priming on different traditional pacing parameters. These findings are not consistent with previous studies on paced mating behavior in female rats that showed that progesterone priming improves CRL compared to EB priming alone (Coopersmith, Candurra, and Erskine 1996; Erskine 1985; Erskine 1992; Brandling-Bennett, Blasberg, and Clark 1999; Chan et al. 2011). The reason for the discrepancy between our study and previous studies could be the cut-off point of 5 seconds that we used to mark an exit, in contrast to an open ending in other studies. This means an escape is considered as an exit only if the female runs from the male to the female compartment within 5 seconds of receiving a stimulation. In most cases in our study, the females did not exit after receiving stimulations, but rather continued copulating or took a pause in the male compartment, and this resulted in fewer exits and, thus, fewer data points for contact-return latency calculations. Because females do not always exit immediately after copulatory stimulations, we have argued that percentage exits and contact return latencies do not fully reflect females’ motivation to continue copulating (reviewed in Heijkoop, Huijgens, and Snoeren 2018; Heijkoop, Huijgens, and Snoeren 2018).

With this study, we hoped to develop an improved assessment tool to study sexual motivation and copulatory rate, and therefore introduced the sexual bouts and time-outs. The observation that female rats copulate in temporal copulatory patterns, structured into sexual bouts and time-outs, is consistent with the copulatory patterns found in male rats (Huijgens et al. 2021). As can be expected, rats with higher levels of receptivity, such as our fully-primed females, consistently copulated with more sexual bouts and shorter time-outs compared to low receptive females. The question that then arises is what do these bouts and time-outs exactly represent? The time-outs are probably the easiest to explain, as they could be considered a measure of motivation to continue copulation. Low receptive female rats are thought to have lower levels of motivation, which is also reflected in our data by having longer time-out durations. Similarly, the fact that the duration of time-outs is related to the type of stimulation the female received in the preceding bout, with an ejaculation lengthening the time-out duration, reflects the concept that higher intensities of stimulations require longer pauses before continuing copulation.

Sexual bouts, on the other hand, are more dependent on the females’ paracopulatory behaviors, and could rather reflect the females’ copulatory intensity. A traditional and comparable parameter in male rats, copulatory rate, is often interpreted as copulatory speed. We previously argued how mount bouts are a better method to describe the copulatory intensity in males than copulatory rate (reviewed in (Heijkoop, Huijgens, and Snoeren 2018)), strengthening the concept of sexual bouts as reflection of the females’ copulatory intensity. Our data shows that sexually experienced females with low receptivity (sub-primed) copulate with fewer, instead of shorter bouts, compared to normal receptive females in a sexual experienced state. This suggests that the number of sexual bouts, rather than the duration of sexual bouts, reflect this copulatory intensity. Although it is hard to define what sexual ‘efficiency’ means for a female rat, the fact that fully-primed females, in contrast to sub-primed females, copulate with more sexual bouts and shorter time-outs, let us conclude that this copulatory patterns should be considered optimal pacing strategy of female rats.

Finally, our data revealed that sexual experience does not strongly affect female sexual behavior, but rather increases the received copulations. Previous research has suggested that sexual experience facilitates sexual performance in female rats (Meerts, Park, and Sekhawat 2016; Meerts et al. 2014; Meerts, Strnad, and Schairer 2015; Blaustein et al. 2009). The reported effects are mainly visible in measures like increased number of received copulations and faster contact-return latencies, but also increased paracopulatory behaviors. Based on this, we hypothesized that sexual experience would modify female rats’ temporal copulatory patterning by reducing time-out duration. Interestingly, although we did not confirm the effects of experience on contact-return latencies or number of paracopulatory behaviors, we observed that sexual experience, in normally functioning fully-primed rats, did reduce the sexual bout durations without affecting the number of sexual bouts or time-out durations. In combination with the observation that the number of paracopulatory behaviors remained stable, this implies that the paracopulatory behaviors are displayed in a shorter period when performed. As such, we found that fully-primed females exhibited significantly more sexual bouts consisting of a single paracopulatory behavior in Cop 6 compared to Cop 1 and Cop 2 (data not shown). Our findings suggest that sexual experience does modify the temporal copulatory patterning of female rats, but the effects are rather small.

## 4 Experiment part 2

In the second part of the experiment, we intended to usefulness of the new behavioral analysis with sexual bouts and time-outs by manipulating the temporal copulatory patterning on female rats. We used (R)-(+)-8-OH-DPAT (DPAT), a serotonin-1A receptor agonist known to inhibit female sexual behavior (Uphouse and Wolf 2004; Kishitake and Yamanouchi 2003; Mendelson and Gorzalka 1986; Snoeren et al. 2010; Snoeren, Refsgaard, et al. 2011; Snoeren et al. 2014), to study the effects on female temporal copulatory patterns in both sub- and fully-primed rats. This allowed us to assess the usefulness of the new behavioral assessment tool and help us interpret the sexual bouts and time-outs.

### 4.1 Results

First, we confirmed our previously reported finding regarding the differences between sub- and fully-primed females in the vehicle (VEH) treated rats. The details of these results can be found in Suppl. Table 4. Here we will continue focusing on the differences in effects of (R)-(+)-8-OH-DPAT (DPAT) treatment.

Our data showed that (R)-(+)-8-OH-DPAT did not affect the time spent in the male compartment of sub- and fully-primed females (Fig. 3A). However, (R)-(+)-8-OH-DPAT significantly reduced the number of paracopulatory behaviors in both hormone groups compared to vehicle treatment (Fig. 3B). Additionally, (R)-(+)-8-OH-DPAT administration led to a decrease in both the lordosis quotient (LQ) and lordosis score (LS) in sub-primed and fully-primed females, relative to vehicle treatment (Fig. 3C/D). Moreover, (R)-(+)-8-OH-DPAT treatment resulted in a reduction in the number of copulatory events, including mounts, intromissions, and ejaculations (Fig. 3E, Suppl. Table 4).

**Figure 3.**
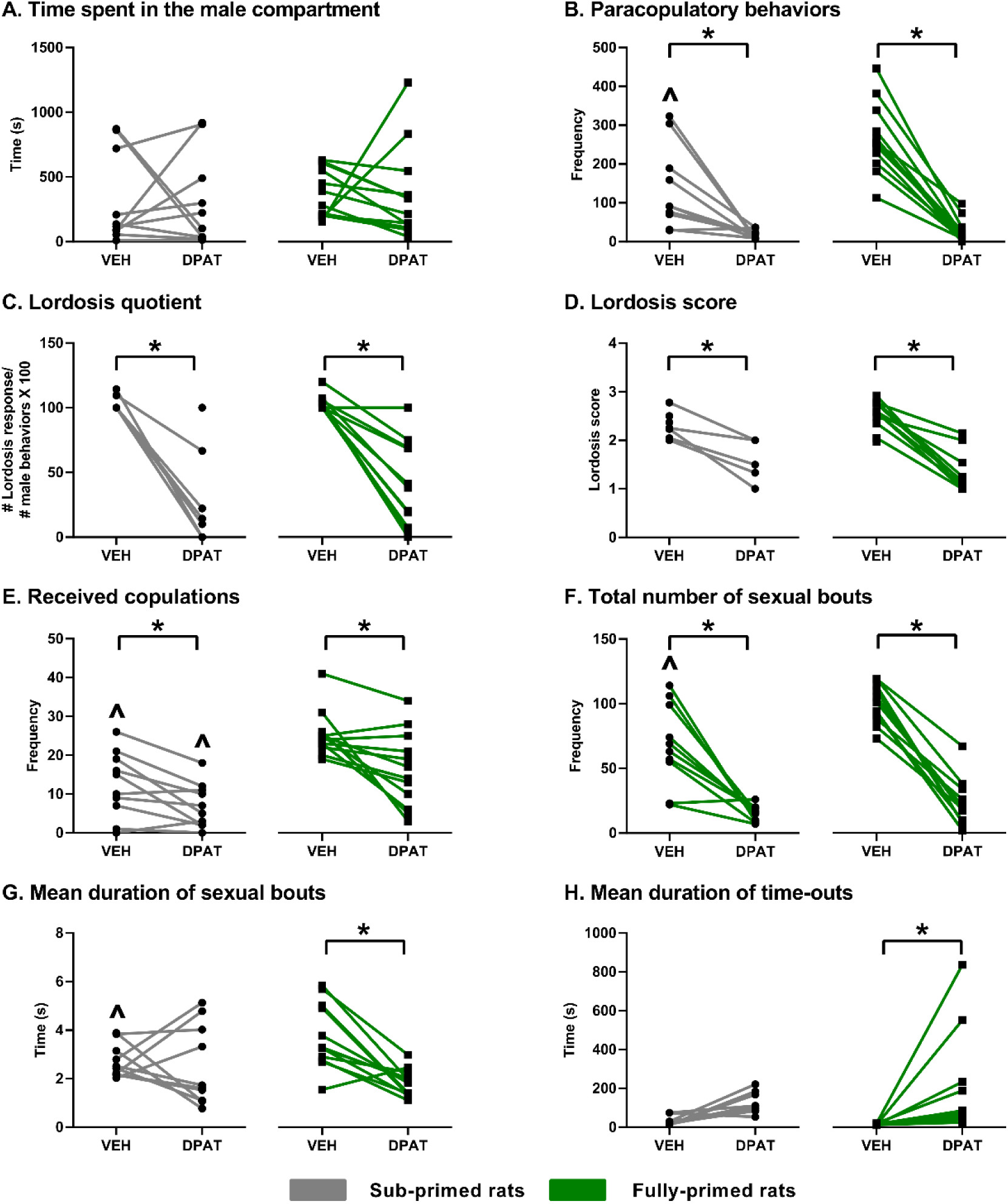
Effects of 0.1mg/kg of (R)-(+)-8-OH-DPAT on female sexual behavior. (A) Time spent in the male compartment (in seconds), (B) Total number of paracopulatory behaviors, (C) Lordosis quotient, (D) Lordosis score, (E) Total number of received copulations, (F) Total number of sexual bouts, (G) Mean duration of sexual bouts (in seconds), (H) Mean duration of time outs (in seconds). **All panels:** Data is shown for sub- and fully-primed female Wistar rats with individual data points, with the lines connecting each rat for both treatments. \**p* < 0.05 significantly different between tests, ^ʌ^*p* < 0.05 is significantly different between groups (sub- vs fully-primed). VEH = vehicle, DPAT = (R)-(+)-8-OH-DPAT.

When looking at the traditional pacing parameters, we found that (R)-(+)-8-OH-DPAT increased the percentage of exits after mounts in sub-primed, but not fully-primed females (Suppl. Table 5). No differences in percentage of exits after intromissions or ejaculations were found between vehicle and (R)-(+)-8-OH-DPAT in either sub- or fully-primed females (Suppl. Table 5). (R)-(+)-8-OH-DPAT did not affect contact return latencies after intromissions and ejaculations, but did increase the CRL after mount in fully-, but not sub-primed rats (Suppl. Table 4).

Next, we investigated whether (R)-(+)-8-OH-DPAT treatment affects the temporal copulatory patterns divided in sexual bouts and time-outs. As expected, our data revealed that (R)-(+)-8-OH-DPAT administration caused a reduction in number of sexual bouts in both sub- and fully-primed females compared to vehicle (Fig. 3F). Additionally, (R)-(+)-8-OH-DPAT shortened the duration of sexual bouts in fully-primed females, but not in sub-primed rats (Fig. 3G). Simultaneously, (R)-(+)-8-OH-DPAT treatment led to a significant increase in the mean duration of time-outs in fully-primed, but not sub-primed, rats (Fig. 3H).

### 4.2 Discussion Experiment part 2

In summary, the results of this 2^nd^ part of the study showed that the temporal copulatory patterns can indeed be manipulated by a sexual inhibiting drug treatment. While (R)-(+)8-OH-DPAT reduced the number of sexual bouts in both sub-and fully-primed females, it only shortened the duration of these bouts and the subsequent time-outs in only fully-primed rats. Furthermore, (R)-(+)-8-OH-DPAT had an inhibiting effect on the number of paracopulatory behaviors, LQ, and LS in both sub- and fully-primed females without clearly affecting traditional pacing parameters.

These results suggest that the serotonin 1A receptor agonist inhibits female sexual behavior by impacting both sexual motivation (as reflected in the duration of time-outs) and copulatory intensity (as reflected in the number and duration of sexual bouts). Our findings are consistent with previous studies, which have also shown that (R)-(+)-8-OH-DPAT inhibits paracopulatory and lordosis behaviors in sub- and fully-primed rats (Snoeren et al. 2010; Snoeren, Refsgaard, et al. 2011; Uphouse, Caldarola-Pastuszka, and Montanez 1992; Ahlenius, Larsson, and Fernandez-Guasti 1989; Kishitake and Yamanouchi 2003; Olivier et al. 2011). The lack of significant changes in traditional pacing parameters, such as exits and contact-return latency, is unsurprising given the limited number of data points. Although some effects on pacing were observed, such as an increase in exits after mounts in sub-primed females and a change in contact-return latency in fully-primed females, these findings were not consistent or corroborated by other metrics.

Overall, this reinforces our view that traditional pacing parameters, such as percentage of exits and contact-return latencies, may be less useful for assessing pacing behavior in female rats. In contrast, our new behavioral paradigm, which divides copulatory behavior into sexual bouts and time-outs, offers a more functional explanation of female sexual performance. Notably, fully-primed rats were more affected by (R)-(+)-8-OH-DPAT administration than sub-primed rats, which may be due to differences in underlying mechanisms or a ceiling effects, given the already low performance of sub-primed females. While research is needed to explore the potential biological underpinnings of these differences, the ability to manipulate new behavioral parameters such as sexual bouts and time-outs, and also obtain subtle differences between treatment groups, underscores the usefulness of this paradigm for advancing our understanding of female rat sexual behavior.

## 5 Experiment part 3

In the final part of the experiment, we aimed to investigate whether a paced mating condition would still have added value over a non-paced condition if this new assessment tool is used for studying female sexual behavior. We therefore compared the temporal copulatory patterns of female rats in a paced mating (PM) vs. non-paced mating (NPM) set-up (Figure 1A).

### 5.1 Results Experiment part 3

Again, we confirmed that sub-primed females show lower levels of sexual activity than fully-primed females. This effect was often found in both non-paced and paced mating settings. The details on this data can be found in Suppl. Table 6, and we only report the relevant findings comparing non-paced to paced mating in this section.

Our data revealed that non-paced mating females displayed more paracopulatory behaviors than paced mating females when fully-primed (Fig. 4A). The latency to start performing paracopulatory behaviors, on the other hand, was not different (Suppl. Table 7). While no differences were found of pacing conditions on lordosis quotient (Fig. 4B), sub-primed females showed a lower lordosis score in the non-paced setting compared to the paced mating set-up (Fig. 4C). Fully-primed females, however showed a similar lordosis score in non-paced and paced mating cages. Furthermore, females received more male copulations in the non-paced mating setting than paced mating set-up (Fig. 4D), especially the sub-primed females.

Since the parameters exits and contact return latencies are not available in non-paced mating -tested rats, we were not able to assess these, but a detailed analysis of sexual bouts and time-outs revealed an increase in the number of sexual bouts in sub-primed females tested in non-paced vs paced mating rats (Fig. 4E). Fully-primed rats, on the other hand, performed the same amount of sexual bouts under paced and non-paced mating conditions. The pacing conditions did not have any effect on the mean duration of the sexual bouts (Fig. 4F), but sub-primed (but not fully-primed) females did have shorter time-outs in the non-paced vs paced mating setting (Fig. 4G).

### 5.2 Discussion Experiment part 2

In summary, our findings show that the mating conditions have a small effect on female sexual performance. In fully-primed females, non-paced mating conditions resulted solely in more paracopulatory behaviors compared to paced mating. Sub-primed females, on the other hand, showed lower LS, but received more copulations in the non-paced mating setting. While no differences were found in fully-primed females on the number and duration of sexual bouts and time-outs, sub-primed females mated with more sexual bouts and shorter time-outs in the non-paced mating vs paced mating set-up.

Our findings are opposite to Coopersmith et al., who showed that female rats tested in a paced mated set-up displayed more paracopulatory behaviors than in a non-paced mating set-up (Coopersmith, Candurra, and Erskine 1996). Another study reported no difference between the total number of paracopulatory behaviors between female rats tested in a paced and non-paced mating set-up (Hernández-Munive et al. 2018).

Overall, this suggests that the conditions in which female rats copulate might not be so relevant when they are normally receptive (fully-primed) but could matter with regard to copulatory intensity and motivation to continue copulation when females are in a low status of receptivity (sub-primed). This could indicate that the presence of, and potentially the extra pressure from, the male could stimulate the female in increasing her copulatory intensity. It should be mentioned, though, that while they can escape in a paced mating setting, female rats in a non-paced mating set-up instead show sexual rejection behaviors. Just as other studies have found (Coopersmith, Candurra, and Erskine 1996; Arzate et al. 2011; Hernández-Munive et al. 2018; Nyuyki et al. 2011), we observed significantly longer sexual rejection behavior episodes (such as fighting, boxing, and kicking) in female rats tested in the non-paced mated set-up (data not shown). This confirms that non-paced mating may not be more engaging but rather more aversive and less rewarding compared to paced mating. The increase in paracopulatory behaviors in fully-primed rats, and the increase in sexual bouts in the sub-primed rats might instead reflect an induced response to the male behavior rather than a voluntary behavior. Altogether, this strengthens the recommendation that paced mating paradigms should be used to study female sexual behavior, as it will lead to a more comprehensive understanding of female sexual behavior and motivation from the female’s willingness to participate in copulatory activity. Moreover, by observing females in a more naturalistic setting, in which they can pace their sexual encounters, we may uncover behavioral consequences, and potentially underlying neural mechanisms, that were previously overlooked or inconclusive in non-paced paradigms.

**Figure 4.**
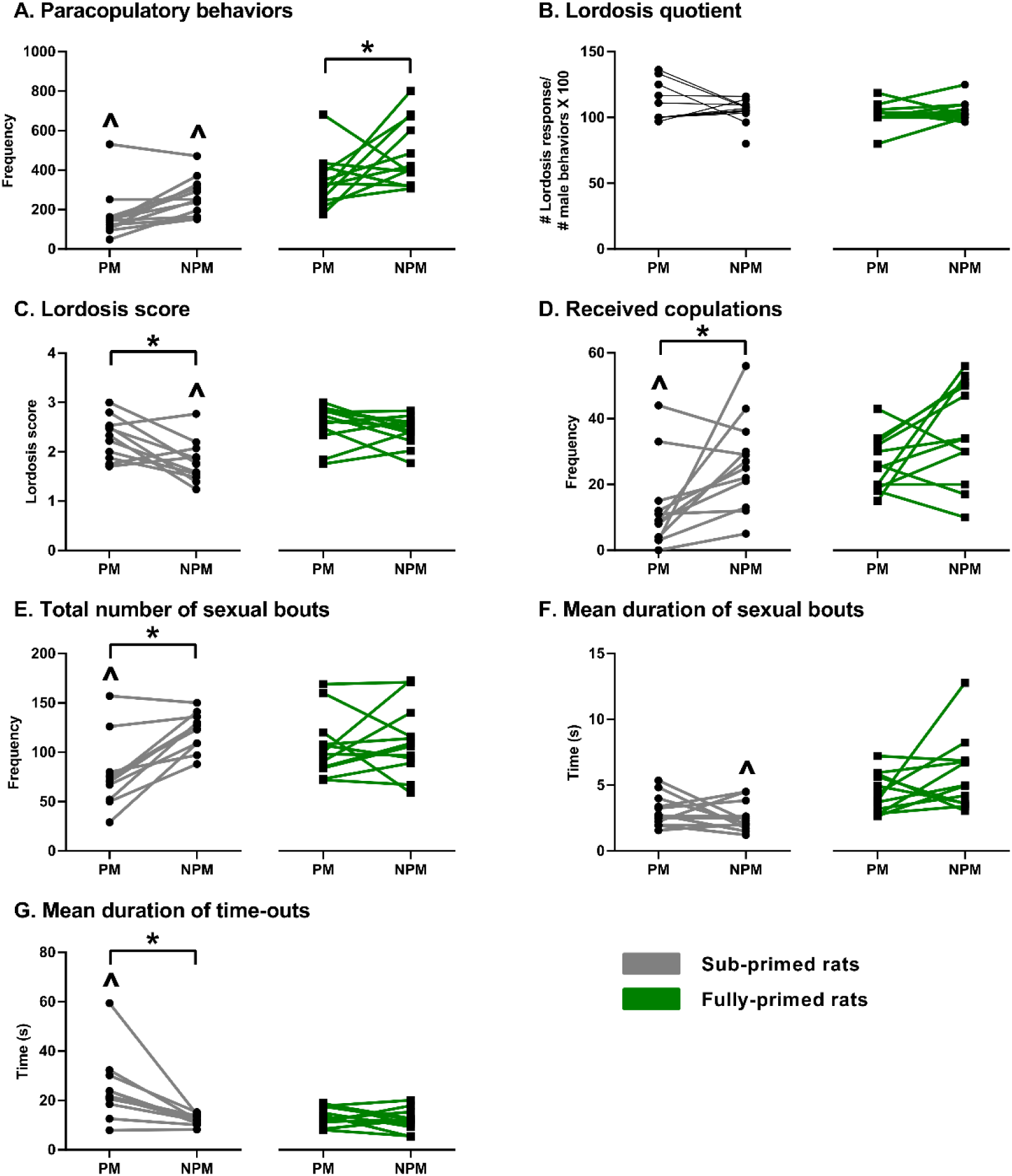
Female sexual behavior in a paced and non-paced mating set-up. (A) Total number of paracopulatory behavior, (B) Lordosis quotient, (C) Lordosis score, (D) Total number of received copulations, (E) Total number of sexual bouts, (F) Mean duration of sexual bouts (in seconds), (G) Mean duration of time outs (in seconds). **All panels:** Data is shown for sub- and fully-primed female Wistar rats with individual data points, with the lines connecting each rat for both Cop tests in different set-up., \**p* < 0.05 is significantly different between tests, ^ʌ^*p* < 0.05 is significantly different between groups (sub- vs fully-primed). PM = paced mating, NPM = non-paced mating.

## 6. Conclusion

Overall, the goal of the study was to develop a new behavioral assessment tool to study temporal copulatory patterns in female rats in more detail. We found that female rats copulate in patterns of sexual bouts and time-outs. While the duration of the sexual bouts solely depends on the female paracopulatory behaviors, the time-out duration is also related to the male’s copulatory stimulation received in the preceding bout. By using low (sub-primed) and normal (fully-primed) receptive females, we determined that higher levels of receptivity result in more sexual bouts and shorter time-outs. This indicated that sexual bouts can be interpreted as measures of copulatory intensity and time-outs as a measure of motivation to continue copulation.

Furthermore, our study showed that sexual experience did not improve the sexual performance of female rats themselves by a large extend, but do result in receiving more mounts, intromissions and ejaculations from the (experienced) male. Finally, we found that the conditions in which female rats copulate (non-paced vs paced mating) might not be so relevant in normal functioning rats, but could matter for females in a low receptive state. Still, sexual rejection is the main focus, paced mating conditions are recommended to study female sexual behavior.

## Author contributions

JCO: Experimental design, data gathering, behavioral annotation, data curation, analysis, writing – original draft.

RH: Experimental design, methodology, data curation, analysis, supervision, writing – review and editing, funding acquisition.

EMSS: Experimental design, methodology, data curation, Programming/software, analysis, supervision, writing – original draft, funding acquisition.

## Acknowledgments

Financial support was received from Helse Nord (HNF1443-19) and the AKM fund of UiT The Arctic university of Norway. We sincerely appreciate Amalie Hofmeyer, Lorenzo Ragazzi, Carina Sørensen, Ragnhild Osnes, Remi Osnes, and Hanna Johansen for the excellent care of the experimental animals. We also extend our gratitude to Truls Traasdahl and the local workshop for their skillful design and construction of our behavioral boxes. Finally, we would also like to thank Dr. Patty Huijgens for revitalizing the concept of the mount bout.

## Notes

### Competing Interest Statement

The authors have declared no competing interest.

