## Supplementary information for "The temporal copulatory patterns of female rat sexual behavior"

**Supplementary table 1.** Definitions of behavioral parameters used in the experiment.

| Behavioral outcome | Definition |
|---|---|
| Paracopulatory behavior | The total number of darts and hops in both male and female compartments. |
| Paracopulatory behaviors in the female compartment | The total number of darts and hops in the female compartment only. |
| Lordosis responses | Lordosis response was assessed on a 4-point scale (0 – 3), with 0 as no lordosis and 3 as a full lordosis with a hollow back and lifted head of 45° or more. From this, we calculate the lordosis quotient and lordosis score. |
| Lordosis quotient | The number of lordosis responses (score 1-3) divided by the number of received copulations (mounts, intromissions, ejaculations) multiplied by 100%. |
| Lordosis score | Lordosis score (number of lordosis responses with score 1 times 1 + number of lordosis responses with score 2 times 2 + number of lordosis responses with score 3 times 3, divided by total number of received copulations.(Hardy and Debold 1971). |
| Mounts | The total number of received mounts. |
| Intromissions | The total number of received intromissions. |
| Ejaculations | The total number of received ejaculations. |
| Received copulations | The total number of received mounts, intromissions, and ejaculations. |
| Time spent in the male compartment | The total time spent in the male compartment. |
| Percentage of exits after mount, intromission, and ejaculations | The total number of withdrawals to the female compartment after received mount, intromission or ejaculation within 5 seconds, divided by the total number of received mounts, intromissions or ejaculations, multiplied by 100%. |
| Contact return latency (CRL) after mount, intromission, and ejaculation | The mean duration for the females to re-enter the male compartment after she withdraw upon a mount, intromission, or ejaculation. |
| Sexual bouts | The total number of sexual bouts defined as a series of behaviors that begins with either a paracopulatory behavior or lordosis response and continues until a behavior was displayed that was not oriented towards the male. The . last paracopulatory or lordosis behavior marked the end of a sexual bout, and the start of a time-out. |

| Behavioral outcome | Definition |
|---|---|
| Mean duration sexual bouts | Mean duration of all sexual bouts, calculated from the first paracopulatory or lordosis behavior in a sexual bout until the last behavior within the sexual bout. |
| Mean duration time-outs | Mean duration of all time-outs, defining a time-out as the time from the end of a sexual bout to the start of the next sexual bout. |
| Paracopulatory behavior in a sexual bout | Mean number of paracopulatory behaviors within a sexual bout that contains at least more than 2 paracopulatory or lordosis behaviors. |
| Lordosis behavior in a sexual bout | Mean number of lordosis behaviors within a sexual bout that contains at least more than 2 paracopulatory or lordosis behaviors. |
| Copulations in a sexual bout | Mean number of mounts and intromissions within a sexual bout that contains at least more than 2 paracopulatory behaviors or lordoses. |

**Supplementary table 2.** Results of the statistical mixed models test in Experiment part 1.

| Parameter | N | Statistics | F value or H value | Post hoc comparison |
|---|---|---|---|---|
| Time spent in male compartment | N = 12 each | Mixed model | Treatment:<br>F(1, 22.000) = 0.006,<br>p = 0.937<br>Experiment:<br>F(3,66) = 31.273,<br>p < 0.001<br>Treatment x Experiment:<br>F(3, 66) = 0.989<br>p = 0.403 | Effect of treatment<br>Cop 1: SP vs FP, p = 0.262<br>Cop 2: SP vs FP, p = 0.475<br>Cop 4: SP vs FP, p = 0.541<br>Cop 6: SP vs FP, p = 0.392<br>Effect of experiment<br>Fully-primed:<br>Cop 1 vs Cop 2, p = 0.001,<br>Cop 1 vs Cop 4, p < 0.001,<br>Cop 1 vs Cop 6, p < 0.001,<br>Cop 2 vs Cop 4, p = 0.190,<br>Cop 2 vs Cop 6, p = 1.000,<br>Cop 4 vs Cop 6, p = 0.231<br>Sub-primed:<br>Cop 1 vs Cop 2, p < 0.001,<br>Cop 1 vs Cop 4, p < 0.001,<br>Cop 1 vs Cop 6, p < 0.001,<br>Cop 2 vs Cop 4, p = 1.000,<br>Cop 2 vs Cop 6, p = 1.000,<br>Cop 4 vs Cop 6, p = 1.000 |
| Total Paracopulatory behavior | N = 12 each | Mixed model | Treatment:<br>F(1,22) = 36.537,<br>p < 0.001<br>Experiment:<br>F(3, 66)= 3.326,<br>p = 0.025<br>Treatment x Experiment:<br>F(3, 66) = 0.900<br>p = 0.446 | Effect of treatment<br>Cop 1: SP vs FP, p < 0.001<br>Cop 2: SP vs FP, p < 0.001<br>Cop 4: SP vs FP, p < 0.001<br>Cop 6: SP vs FP, p < 0.001<br>Effect of experiment<br>Fully-primed<br>Cop 1 vs Cop 2, p = 1.000,<br>Cop 1 vs Cop 4, p = 0.416,<br>Cop 1 vs Cop 6, p = 1.000,<br>Cop 2 vs Cop 4, p = 1.000,<br>Cop 2 vs Cop 6, p = 1.000,<br>Cop 4 vs Cop 6, p = 0.122,<br>Sub-primed:<br>Cop 1 vs Cop 2, p = 0.996,<br>Cop 1 vs Cop 4, p = 1.000,<br>Cop 1 vs Cop 6, p = 0.482,<br>Cop 2 vs Cop 4, p = 0.592,<br>Cop 2 vs Cop 6, p = 1.000,<br>Cop 4 vs Cop 6, p = 0.266 |
| Paracopulatory in the female compartment | N = 12 each | Mixed model | Treatment:<br>F(1, 22.000) = 26.541,<br>p < 0.001<br>Experiment:<br>F(3,66) = 13.754,<br>p < 0.001<br>Treatment x Experiment:<br>F(3, 66) = 4.107,<br>p = 0.010 | Effect of treatment<br>Cop 1: SP vs FP, p = 0.011,<br>Cop 2: SP vs FP, p = 0.017,<br>Cop 4: SP vs FP, p < 0.001,<br>Cop 6: SP vs FP, p = 0.010,<br>Effect of experiment<br>Fully-primed:<br>Cop 1 vs Cop 2, p = 0.002,<br>Cop 1 vs Cop 4, p < 0.001,<br>Cop 1 vs Cop 6, p = 0.212,<br>Cop 2 vs Cop 4, p = 0.179,<br>Cop 2 vs Cop 6, p = 0.713,<br>Cop 4 vs Cop 6, p = 0.002<br>Sub-primed:<br>Cop 1 vs Cop 2, p = 0.001, |

| | | | | |
|---|---|---|---|---|
| | | | | Cop 1 vs Cop 4, p = 0.315,<br>Cop 1 vs Cop 6, p = 0.218,<br>Cop 2 vs Cop 4, p = 0.310,<br>Cop 2 vs Cop 6, p = 0.441,<br>Cop 4 vs Cop 6, p = 1.000 |
| Lordosis quotient | N = 12 each | Mixed model | Treatment:<br>F(1, 28.402) = 1.967,<br>p = 0.172<br>Experiment:<br>F(3,59.838) = 2.676,<br>p = 0.055<br>Treatment x Experiment:<br>F(3, 59.838) = 3.629,<br>p = 0.018 | Effect of treatment<br>Cop 1: SP vs FP, p = 0.005,<br>Cop 2: SP vs FP, p = 0.536,<br>Cop 4: SP vs FP, p = 0.192,<br>Cop 6: SP vs FP, p = 0.512,<br>Effect of experiment<br>Fully-primed:<br>Cop 1 vs Cop 2, p < 0.001, Cop 1 vs Cop 4, p < 0.001, Cop 1 vs Cop 6, p < 0.001, Cop 2 vs Cop 4, p = 1.000, Cop 2 vs Cop 6, p = 1.000,<br>Cop 4 vs Cop 6, p = 1.000,<br>Sub-primed:<br>Cop 1 vs Cop 2, p = 1.000,<br>Cop 1 vs Cop 4, p = 1.000,<br>Cop 1 vs Cop 6, p = 1.000,<br>Cop 2 vs Cop 4, p = 1.000,<br>Cop 2 vs Cop 6, p = 1.000,<br>Cop 4 vs Cop 6, p = 1.000 |
| Lordosis score | N = 12 each | Mixed model | Treatment:<br>F(1, 18.939) = 2.132,<br>p = 0.161<br>Experiment:<br>F(3,56.641) = 4.258,<br>p = 0.009<br>Treatment x Experiment:<br>F(3, 56.641) = 0.452,<br>p = 0.717 | Effect of treatment<br>Cop 1: SP vs FP, p = 0.259,<br>Cop 2: SP vs FP, p = 0.320,<br>Cop 4: SP vs FP, p = 0.818,<br>Cop 6: SP vs FP, p = 0.115,<br>Effect of experiment<br>Fully-primed:<br>Cop 1 vs Cop 2, p = 1.000, Cop 1 vs Cop 4, p = 0.956, Cop 1 vs Cop 6, p = 0.037, Cop 2 vs Cop 4, p = 1.000, Cop 2 vs Cop 6, p = 0.716,<br>Cop 4 vs Cop 6, p = 0.956,<br>Sub-primed:<br>Cop 1 vs Cop 2, p = 1.000, Cop 1 vs Cop 4, p = 0.171, Cop 1 vs Cop 6, p = 0.258, Cop 2 vs Cop 4, p = 1.000,<br>Cop 2 vs Cop 6, p = 1.000, Cop 4 vs Cop 6, p = 1.000 |
| Received copulations | N = 12 each | Mixed model | Treatment:<br>F(1,22) = 22.649,<br>p < 0.001<br>Experiment:<br>F(3, 66)= 11.028,<br>p < 0.001<br>Treatment x Experiment:<br>F(3, 66) = 0.384,<br>p = 0.765 | Effect of treatment<br>Cop 1: SP vs FP, p < 0.001,<br>Cop 2: SP vs FP, p = 0.002,<br>Cop 4: SP vs FP, p = 0.012,<br>Cop 6: SP vs FP, p < 0.001,<br>Effect of experiment<br>Fully-primed:<br>Cop 1 vs Cop 2, p = 0.018, Cop 1 vs Cop 4, p = 1.000, Cop 1 vs Cop 6, p = 0.006, Cop 2 vs Cop 4, p = 0.306, Cop 2 vs Cop 6, p = 1.000,<br>Cop 4 vs Cop 6, p = 0.138,<br>Sub-primed:<br>Cop 1 vs Cop 2, p = 0.002, Cop 1 vs Cop 4, p = 0.087, Cop 1 vs Cop 6, p = 0.003, Cop 2 vs Cop 4, p = |

| | | | | |
|---|---|---|---|---|
| | | | | 1.000, Cop 2 vs Cop 6, p = 1.000, Cop 4 vs Cop 6, p = 1.000 |
| Latency to 1st paracopulatory | N = 12 each | Mixed model | Treatment: $F(1, 22.000) = 11.877$, $p = 0.002$<br>Experiment: $F(3,66) = 19.118$, $p < 0.001$<br>Treatment x Experiment: $F(3, 66) = 5.915$, $p = 0.001$ | Effect of treatment<br>Cop 1: SP vs FP, p < 0.001,<br>Cop 2: SP vs FP, p = 0.181,<br>Cop 4: SP vs FP, p = 0.331,<br>Cop 6: SP vs FP, p = 0.802,<br>Effect of experiment<br>Fully-primed:<br>Cop 1 vs Cop 2, p = 0.531, Cop 1 vs Cop 4, p = 0.510, Cop 1 vs Cop 6, p = 0.187, Cop 2 vs Cop 4, p = 1.000, Cop 2 vs Cop 6, p = 1.000, Cop 4 vs Cop 6, p = 1.000,<br>Sub-primed:<br>Cop 1 vs Cop 2, p < 0.001, Cop 1 vs Cop 4, p < 0.001, Cop 1 vs Cop 6, p < 0.001, Cop 2 vs Cop 4, p = 1.000, Cop 2 vs Cop 6, p = 0.640, Cop 4 vs Cop 6, p = 1.000 |
| Mounts | N = 12 each | Mixed model | Treatment: $F(1, 22.000) = 16.732$, $p < 0.001$<br>Experiment: $F(3,66) = 2.335$, $p = 0.082$<br>Treatment x Experiment: $F(3, 66) = 1.026$, $p = 0.387$ | Effect of treatment<br>Cop 1: SP vs FP, p = 0.005,<br>Cop 2: SP vs FP, p = 0.047,<br>Cop 4: SP vs FP, p = 0.341,<br>Cop 6: SP vs FP, p = 0.002,<br>Effect of experiment<br>Fully-primed:<br>Cop 1 vs Cop 2, p = 1.000, Cop 1 vs Cop 4, p = 1.000, Cop 1 vs Cop 6, p = 0.736,<br>Cop 2 vs Cop 4, p = 0.665, Cop 2 vs Cop 6, p = 1.000,<br>Cop 4 vs Cop 6, p = 0.044,<br>Sub-primed:<br>Cop 1 vs Cop 2, p = 1.000, Cop 1 vs Cop 4, p = 1.000, Cop 1 vs Cop 6, p = 1.000,<br>Cop 2 vs Cop 4, p = 1.000, Cop 2 vs Cop 6, p = 1.000,<br>Cop 4 vs Cop 6, p = 1.000 |
| Intromissions | N = 12 each | Mixed model | Treatment: $F(1, 22.000) = 16.582$, $p < 0.001$<br>Experiment: $F(3,66) = 13.231$, $p < 0.001$<br>Treatment x Experiment: $F(3, 66) = 0.217$, $P = 0.884$ | Effect of treatment<br>Cop 1: SP vs FP, p < 0.001,<br>Cop 2: SP vs FP, p = 0.003,<br>Cop 4: SP vs FP, p = 0.011,<br>Cop 6: SP vs FP, p = 0.005,<br>Effect of experiment<br>Fully-primed:<br>Cop 1 vs Cop 2, p = 0.002, Cop 1 vs Cop 4, p = 0.526, Cop 1 vs Cop 6, p = 0.008.<br>Cop 2 vs Cop 4, p = 0.283, Cop 2 vs Cop 6, p = 1.000,<br>Cop 4 vs Cop 6, p = 0.673,<br>Sub-primed: |

| | | | | |
|---|---|---|---|---|
| | | | | Cop 1 vs Cop 2, p < 0.001, Cop 1 vs Cop 4, p = 0.035, Cop 1 vs Cop 6, p < 0.001,<br>Cop 2 vs Cop 4, p = 1.000, Cop 2 vs Cop 6, p = 1.000,<br>Cop 4 vs Cop 6, p = 1.000 |
| Ejaculations | N = 12 each | Mixed model | Treatment:<br>F(1, 22.000) = 7.508,<br>p = 0.012<br>Experiment:<br>F(3,66) = 12.600,<br>p < 0.001<br>Treatment x Experiment: F(3, 66) = 0.879,<br>p = 0.457 | Effect of treatment<br>Cop 1: SP vs FP, p = 0.246,<br>Cop 2: SP vs FP, p = 0.099,<br>Cop 4: SP vs FP, p = 0.004,<br>Cop 6: SP vs FP, p = 0.099,<br>Effect of experiment<br>Fully-primed:<br>Cop 1 vs Cop 2, p = 0.002, Cop 1 vs Cop 4, p < 0.001, Cop 1 vs Cop 6, p = 0.001,<br>Cop 2 vs Cop 4, p = 1.000, Cop 2 vs Cop 6, p = 1.000,<br>Cop 4 vs Cop 6, p = 0.673,<br>Sub-primed:<br>Cop 1 vs Cop 2, p = 0.013, Cop 1 vs Cop 4, p = 0.120, Cop 1 vs Cop 6, p = 0.007,<br>Cop 2 vs Cop 4, p = 1.000, Cop 2 vs Cop 6, p = 1.000,<br>Cop 4 vs Cop 6, p = 1.000 |
| Latency to 1st ejaculation | N = 12 each | Mixed model | Treatment:<br>F(1, 22.000) = 18.159,<br>p < 0.001<br>Experiment:<br>F(3,66) = 13.868,<br>p < 0.001<br>Treatment x Experiment: F(3, 66) = 0.648,<br>p = 0.587 | Effect of treatment<br>Cop 1: SP vs FP, p = 0.043<br>Cop 2: SP vs FP, p = 0.024<br>Cop 4: SP vs FP, p < 0.001<br>Cop 6: SP vs FP, p = 0.013<br>Effect of experiment<br>Fully-primed:<br>Cop 1 vs Cop 2, p = 0.001, Cop 1 vs Cop 4, p < 0.001, Cop 1 vs Cop 6, p = 0.001,<br>Cop 2 vs Cop 4, p = 1.000, Cop 2 vs Cop 6, p = 1.000,<br>Cop 4 vs Cop 6, p = 1.000,<br>Sub-primed:<br>Cop 1 vs Cop 2, p = 0.002, Cop 1 vs Cop 4, p = 0.050, Cop 1 vs Cop 6, p = 0.007,<br>Cop 2 vs Cop 4, p = 1.000, Cop 2 vs Cop 6, p = 1.000,<br>Cop 4 vs Cop 6, p = 1.000 |
| Percentage of exits after mounts | N = 12 each | Mixed model | Treatment:<br>F(1, 19.894) = 1.550,<br>p = 0.228<br>Experiment:<br>F(3,46.286) = 1.274,<br>p = 0.294<br>Treatment x Experiment: F(3, 46.286) = 0.443,<br>P = 0.723 | Effect of treatment<br>Cop 1: SP vs FP, p = 0.346,<br>Cop 2: SP vs FP, p = 0.122,<br>Cop 4: SP vs FP, p = 0.661,<br>Cop 6: SP vs FP, p = 0.433,<br>Effect of experiment<br>Fully-primed: |

| | | | | |
|---|---|---|---|---|
| | | | | Cop 1 vs Cop 2, p = 0.829, Cop 1 vs Cop 4, p = 0.499, Cop 1 vs Cop 6, p = 1.000,<br>Cop 2 vs Cop 4, p = 1.000,<br>Cop 2 vs Cop 6, p = 1.000,<br>Cop 4 vs Cop 6, p = 1.000,<br>Sub-primed:<br>Cop 1 vs Cop 2, p = 1.000,<br>Cop 1 vs Cop 4, p = 1.000,<br>Cop 1 vs Cop 6, p = 1.000,<br>Cop 2 vs Cop 4, p = 1.000,<br>Cop 2 vs Cop 6, p = 1.000,<br>Cop 4 vs Cop 6, p = 1.000 |
| Percentage of exits after intromissions | N = 12 each | Mixed model | Treatment:<br>F(1, 31.068) = 0.416,<br>p = 0.524<br>Experiment:<br>F(3,48.449) = 12.825,<br>p < 0.001<br>Treatment x Experiment: F(3, 48.449) = 1.166,<br>p = 0.332 | Effect of treatment<br>Cop 1: SP vs FP, p = 0.694,<br>Cop 2: SP vs FP, p = 0.598,<br>Cop 4: SP vs FP, p = 0.417,<br>Cop 6: SP vs FP, p = 0.211,<br>Effect of experiment<br>Fully-primed:<br>Cop 1 vs Cop 2, p = 0.173, Cop 1 vs Cop 4, p < 0.001,<br>Cop 1 vs Cop 6, p < 0.001,<br>Cop 2 vs Cop 4, p = 0.052, Cop 2 vs Cop 6, p = 0.208,<br>Cop 4 vs Cop 6, p = 1.000,<br>Sub-primed:<br>Cop 1 vs Cop 2, p = 1.000, Cop 1 vs Cop 4, p = 0.413, Cop 1 vs Cop 6, p = 0.386,<br>Cop 2 vs Cop 4, p < 0.001, Cop 2 vs Cop 6, p < 0.001,<br>Cop 4 vs Cop 6, p = 1.000 |
| Percentage of exits after ejaculations | N = 12 each | Mixed model | Treatment:<br>F(1, 22.152) = 3.312,<br>p = 0.082<br>Experiment:<br>F(3,42.262) = 1.568,<br>p = 0.211<br>Treatment x Experiment: F(3, 48.449) = 0.084,<br>p = 0.920 | Effect of treatment<br>Cop 1: SP vs FP: Not enough data points<br>Cop 2: SP vs FP, p = 0.340<br>Cop 4: SP vs FP, p = 0.429<br>Cop 6: SP vs FP, p = 0.167<br>Effect of experiment<br>Fully-primed:<br>Cop 1 vs Cop 2, p = 1.000,<br>Cop 1 vs Cop 4, p = 1.000,<br>Cop 1 vs Cop 6, p = 1.000,<br>Cop 2 vs Cop 4, p = 1.000, Cop 2 vs Cop 6, p = 0.912,<br>Cop 4 vs Cop 6, p = 1.000,<br>Sub-primed:<br>Cop 1 vs Cop 2: Not enough data points,<br>Cop 1 vs Cop 4: Not enough data points, |

| | | | | |
|---|---|---|---|---|
| | | | | Cop 1 vs Cop 6: Not enough data points,<br>Cop 2 vs Cop 4, p = 1.000, Cop 2 vs Cop 6, p = 0.420,<br>Cop 4 vs Cop 6, p = 1.000 |
| Contact return latency after mounts | N = 12 each | Mixed model | Treatment:<br>F(1, 10.73) = 0.207,<br>p = 0.658<br>Experiment:<br>F(3, 13.59) = 0.476,<br>p = 0.704<br>Treatment* experiment: F(3, 13.59) = 0.595,<br>p = 0.629 | Effect of treatment<br>Cop 1: SP vs FP, p = 0.830,<br>Cop 2: SP vs FP, p = 0.185,<br>Cop 4: SP vs FP, p = 0.659,<br>Cop 6: SP vs FP, p = 0.971,<br>Effect of experiment<br>Fully-primed:<br>Cop 1 vs Cop 2, p = 1.000, Cop 1 vs Cop 4, p = 1.000,<br>Cop 1 vs Cop 6, p = 1.000,<br>Cop 2 vs Cop 4, p = 1.000, Cop 2 vs Cop 6, p = 1.000,<br>Cop 4 vs Cop 6, p = 1.000,<br>Sub-primed:<br>Cop 1 vs Cop 2, p = 1.000,<br>Cop 1 vs Cop 4, p = 1.000,<br>Cop 1 vs Cop 6, p = 1.000,<br>Cop 2 vs Cop 4, p = 1.000, Cop 2 vs Cop 6, p = 1.000,<br>Cop 4 vs Cop 6, p = 1.000 |
| Contact return latency after intromissions | N = 12 each | Mixed model | Treatment:<br>F(1,17.42) = 0.038,<br>p = 0.849<br>Experiment:<br>F(3, 36.05) = 1.968,<br>p = 0.136<br>Treatment* experiment: F(2, 32.53) = 0.889,<br>p = 0.421 | Effect of treatment<br>Cop 1: SP vs FP: Not enough data points,<br>Cop 2: SP vs FP, p = 0.332,<br>Cop 4: SP vs FP, p = 0.946,<br>Cop 6: SP vs FP, p = 0.346,<br>Effect of experiment<br>Fully-primed:<br>Cop 1 vs Cop 2, p = 1.000, Cop 1 vs Cop 4, p = 1.000,<br>Cop 1 vs Cop 6, p = 1.000,<br>Cop 2 vs Cop 4, p = 0.349, Cop 2 vs Cop 6, p = 1.000,<br>Cop 4 vs Cop 6, p = 0.279<br>Sub-primed: '<br>Cop 1 vs Cop 2: Not enough data points,<br>Cop 2 vs Cop 4, p = 0.252, Cop 2 vs Cop 6, p = 1.000,<br>Cop 4 vs Cop 6, p = 0.859 |
| Contact return latency after ejaculations | N = 12 each | Mixed model | Treatment:<br>F(1,9.319) = 1.485,<br>p = 0.253<br>Experiment:<br>F(3, 19.022) = 4.085,<br>p = 0.021<br>Treatment* experiment: F(2, 16.507) = 2.404,<br>p = 0.121. | Effect of treatment<br>Cop 1: SP vs FP: Not enough data points,<br>Cop 2: SP vs FP, p = 0.037,<br>Cop 4: SP vs FP, p = 0.437,<br>Cop 6: SP vs FP, p = 0.576,<br>Effect of experiment<br>Fully-primed: |

| | | | | |
|---|---|---|---|---|
| | | | | Cop 1 vs Cop 2, p = 1.000, Cop 1 vs Cop 4, p = 0.486,<br>Cop 1 vs Cop 6, p = 1.000,<br>Cop 2 vs Cop 4, p = 1.000, Cop 2 vs Cop 6, p = 1.000,<br>Cop 4 vs Cop 6, p = 1.000,<br>Sub-primed:<br>Cop 1 vs Cop 2: Not enough data points,<br>Cop 1 vs Cop 4: Not enough data points,<br>Cop 1 vs Cop 6: Not enough data points,<br>Cop 2 vs Cop 4, p = 0.009, Cop 2 vs Cop 6, p = 0.470,<br>Cop 4 vs Cop 6, p = 0.160 |
| Total number of sexual bouts | N = 12 each | Mixed model | Treatment:<br>F(1, 22) = 12.637,<br>p = 0.002<br>Experiment:<br>F(3,66) = 2.490,<br>p = 0.068<br>Treatment x Experiment: F(3, 66) = 2.586,<br>p = 0.060 | Effect of treatment<br>Cop 1: SP vs FP, p = 0.023,<br>Cop 2: SP vs FP, p = 0.212,<br>Cop 4: SP vs FP, p < 0.001,<br>Cop 6: SP vs FP, p = 0.013,<br>Effect of experiment<br>Fully-primed:<br>Cop 1 vs Cop 2, p = 1.000, Cop 1 vs Cop 4, p = 1.000,<br>Cop 1 vs Cop 6, p = 0.394,<br>Cop 2 vs Cop 4, p = 1.000, Cop 2 vs Cop 6, p = 1.000,<br>Cop 4 vs Cop 6, p = 0.236,<br>Sub-primed:<br>Cop 1 vs Cop 2, p = 1.000, Cop 1 vs Cop 4, p = 1.000, Cop 1 vs Cop 6, p = 0.739,<br>Cop 2 vs Cop 4, p = 0.080, Cop 2 vs Cop 6, p = 1.000,<br>Cop 4 vs Cop 6, p = 0.049 |
| Mean duration sexual bouts | N = 12 each | Mixed model | Treatment:<br>F(1, 22) = 33.890,<br>p < 0.001<br>Experiment:<br>F(3,66) = 4.191,<br>p = 0.009<br>Treatment x Experiment: F(3, 66) = 5.540,<br>p = 0.002 | Effect of treatment<br>Cop 1: SP vs FP, p < 0.001,<br>Cop 2: SP vs FP, p < 0.001,<br>Cop 4: SP vs FP, p < 0.001,<br>Cop 6: SP vs FP, p = 0.066,<br>Effect of experiment<br>Fully-primed:<br>Cop 1 vs Cop 2, p = 1.000, Cop 1 vs Cop 4, p = 0.045,<br>Cop 1 vs Cop 6, p < 0.001,<br>Cop 2 vs Cop 4, p = 0.360, Cop 2 vs Cop 6, p < 0.001,<br>Cop 4 vs Cop 6, p = 0.242,<br>Sub-primed:<br>Cop 1 vs Cop 2, p = 1,000, Cop 1 vs Cop 4, p = 1.000, Cop 1 vs Cop 6, p = 1.000,<br>Cop 2 vs Cop 4, p = 1.000, Cop 2 vs Cop 6, p = 1.000, |

| | | | | |
|---|---|---|---|---|
| | | | | Cop 4 vs Cop 6, p = 1.000 |
| Mean duration time-outs | N = 12 each | Mixed model | Treatment:<br>F(1, 22) = 15.025,<br>p < 0.001<br>Experiment:<br>F(3,66) = 2.957,<br>p = 0.039<br>Treatment x Experiment: F(3, 66) = 3.191,<br>p = 0.029 | Effect of treatment<br>Cop 1: SP vs FP, p = 0.007,<br>Cop 2: SP vs FP, p = 0.046,<br>Cop 4: SP vs FP, p < 0.001,<br>Cop 6: SP vs FP, p = 0.048,<br>Effect of experiment<br>Fully-primed:<br>Cop 1 vs Cop 2, p = 1.000, Cop 1 vs Cop 4, p = 1.000,<br>Cop 1 vs Cop 6, p = 1.000,<br>Cop 2 vs Cop 4, p = 1.000, Cop 2 vs Cop 6, p = 1.000,<br>Cop 4 vs Cop 6, p = 1.000,<br>Sub-primed:<br>Cop 1 vs Cop 2, p = 1,000, Cop 1 vs Cop 4, p = 0.047, Cop 1 vs Cop 6, p = 1.000,<br>Cop 2 vs Cop 4, p = 0.003, Cop 2 vs Cop 6, p = 1.000,<br>Cop 4 vs Cop 6, p = 0.002 |
| Paracopulatory per sexual bout | N = 12 each | Mixed model | Treatment:<br>F(1, 22.000) = 31.924,<br>p < 0.001<br>Experiment:<br>F(3,66) = 3.825,<br>p = 0.014<br>Treatment x Experiment: F(3, 66) = 2.437,<br>p = 0.073 | Effect of treatment<br>Cop 1: SP vs FP, p < 0.001,<br>Cop 2: SP vs FP, p = 0.002,<br>Cop 4: SP vs FP, p = 0.015,<br>Cop 6: SP vs FP, p < 0.001,<br>Effect of experiment<br>Fully-primed:<br>Cop 1 vs Cop 2, p = 0.865, Cop 1 vs Cop 4, p = 0.005, Cop 1 vs Cop 6, p = 1.000,<br>Cop 2 vs Cop 4, p = 0.290, Cop 2 vs Cop 6, p = 1.000,<br>Cop 4 vs Cop 6, p = 0.008,<br>Sub-primed:<br>Cop 1 vs Cop 2, p = 1.000, Cop 1 vs Cop 4, p = 1.000, Cop 1 vs Cop 6, p = 0.662,<br>Cop 2 vs Cop 4, p = 1.000, Cop 2 vs Cop 6, p = 1.000,<br>Cop 4 vs Cop 6, p = 1.000 |
| Lordoses per sexual bout | N = 12 each | Mixed model | Treatment:<br>F(1, 21.757) = 12.465,<br>p = 0.002<br>Experiment:<br>F(3,64.649) = 4.388,<br>p = 0.007<br>Treatment x Experiment: F(3, 64.649) = 1.501,<br>p = 0.223 | Effect of treatment<br>Cop 1: SP vs FP, p = 0.002,<br>Cop 2: SP vs FP, p = 0.108,<br>Cop 4: SP vs FP, p = 0.719,<br>Cop 6: SP vs FP, p = 0.016,<br>Effect of experiment<br>Fully-primed:<br>Cop 1 vs Cop 2, p = 1.000, Cop 1 vs Cop 4, p = 1.000, Cop 1 vs Cop 6, p = 0.248,<br>Cop 2 vs Cop 4, p = 1.000, Cop 2 vs Cop 6, p = 1.000,<br>Cop 4 vs Cop 6, p = 0.515,<br>Sub-primed: |

| | | | | |
|---|---|---|---|---|
| | | | | Cop 1 vs Cop 2, p = 0.098, Cop 1 vs Cop 4, p = 0.014, Cop 1 vs Cop 6, p = 0.038,<br>Cop 2 vs Cop 4, p = 1.000, Cop 2 vs Cop 6, p = 1.000,<br>Cop 4 vs Cop 6, p = 1.000 |
| Copulations per sexual bout | N = 12 each | Mixed model | Treatment:<br>F(1, 21.737) = 12.462,<br>p = 0.002<br>Experiment:<br>F(3,64.583) = 5.578,<br>p = 0.002<br>Treatment x Experiment: F(3, 64.583) = 1.369,<br>p = 0.260 | Effect of treatment<br>Cop 1: SP vs FP, p = 0.002,<br>Cop 2: SP vs FP, p = 0.094,<br>Cop 4: SP vs FP, p = 0.619,<br>Cop 6: SP vs FP, p = 0.013,<br>Effect of experiment<br>Fully-primed:<br>Cop 1 vs Cop 2, p = 1.000, Cop 1 vs Cop 4, p = 1.000, Cop 1 vs Cop 6, p = 0.100,<br>Cop 2 vs Cop 4, p = 1.000, Cop 2 vs Cop 6, p = 1.000,<br>Cop 4 vs Cop 6, p = 0.413,<br>Sub-primed:<br>Cop 1 vs Cop 2, p = 0.062, Cop 1 vs Cop 4, p = 0.010, Cop 1 vs Cop 6, p = 0.018,<br>Cop 2 vs Cop 4, p = 1.000, Cop 2 vs Cop 6, p = 1.000,<br>Cop 4 vs Cop 6, p = 1.000 |
| Sexual bout duration per sexual bout type | N = 7675 | Kruskal-Wallis | H(3) = 1334.288,<br>p < 0.001 | 1-5P vs 6-10P:<br>Z = -28.138, p < 0.001,<br>1-5P vs 11-20P:<br>Z = - 20.787, p < 0.001,<br>1-5P vs >20P:<br>Z = -13.308, p < 0.001,<br>6-10P vs 11-20P:<br>Z = -7.932, p < 0.001,<br>6-10P vs >20P:<br>Z = - 10.597, p < 0.001,<br>11-20P vs >20P:<br>Z = - 7.629, p < 0.001 |
| Sexual bout duration per copulation type | N = 1190 | Kruska-Wallis | H(4) = 128.455,<br>p < 0.001 | M only vs 1I:<br>Z = -4.195, p < 0.001,<br>M only vs 1E:<br>Z = -1.941, p = 0.052,<br>M only vs M+1:<br>Z = -5.358, p < 0.001,<br>M only vs M+I+E:<br>Z = -3.658, p < 0.001,<br>1I vs 1E:<br>Z = -6.279, p < 0.001,<br>1I vs M+I:<br>Z = -9.137, p < 0.001,<br>1I vs M+I+E:<br>Z = -4.000, p < 0.001,<br>M+I vs M+I+E:<br>Z = -3.167, p = 0.002,<br>1E vs M+I:<br>Z = -3.895, p < 0.001,<br>1E vs M+I+E:<br>Z = -3.684, p < 0.001 |

| | | | | |
|---|---|---|---|---|
| Time-out duration per sexual bout type | N = 7587 | Kruskal-Wallis | H(3) = 0.806, p = 0.848 | 1-5P vs 6-10P: Z = - 0.163, p = 0.870, 1-5P vs 11-20P: Z = - 0.192, p = 0.848, 1-5P vs >20P: Z = - 0.870, p = 0.384, 6-10P vs 11-20P: Z = -0.092, p = 0.927, 6-10P vs >20P: Z = - 0.755, p = 0.450, 11-20P vs >20P: Z = - 0.699, p = 0.484, |
| Time-out duration per copulation type | N = 1166 | Kruskal-Wallis | H(4) = 128.455, p < 0.001 | M only vs 1I: Z = -10.042, p < 0.001, M only vs 1E: Z = -13.910, p < 0.001, M only vs M+1: Z = -3.938, p < 0.001, M only vs M+I+E: Z = -3.897, p < 0.001, 1I vs 1E: Z = -14.438, p < 0.001, 1I vs M+I: Z = -2.860, p = 0.004, 1I vs M+I+E: Z = -3.177, p = 0.001, M+I vs M+I+E: Z = -3.351, p < 0.001, 1E vs M+I: Z = -10.315, p < 0.001, 1E vs M+I+E: Z = -0.049, p = 0.961 |

S1. Paracopulatory behaviors in the female compartment

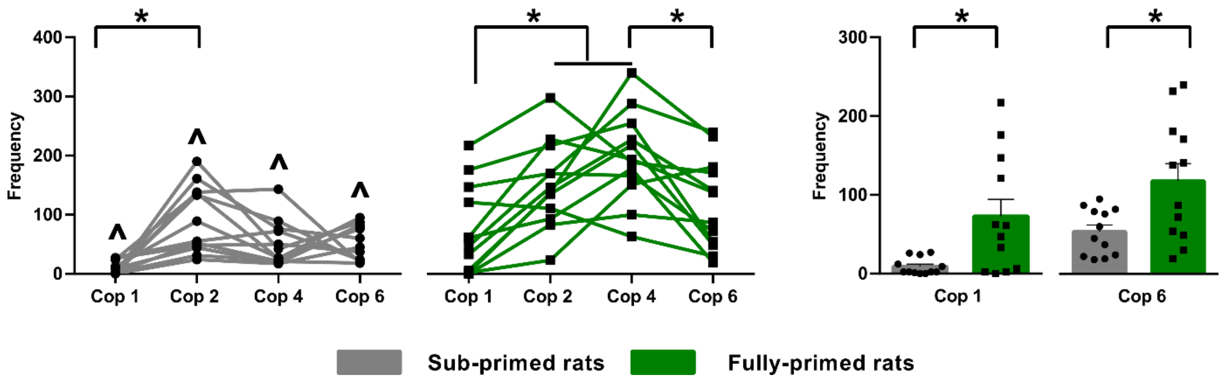


**Supplementary Figure S1.** Sexual experience and hormonal status and the total number of paracopulatory behaviors performed in the female compartment. **Panel to the left**: The data are shown with individual data points, with the lines connecting each rat across different Cop tests; **panel to the right:** The data are shown with individual data points, with the bars representing the mean ± SEM, **All figures**: Data is shown for sub- and fully-primed female Wistar rats. *$p < 0.05$ significantly different between Cop tests, ^$p < 0.05$ is significantly different between groups (sub- vs fully-primed). Cop = Copulation test.

**Supplementary table 3.** Effects of hormonal priming and sexual experience on female sexual behavior.

| **Parameters** | | **Cop 1** | **Cop 2** | **Cop 4** | **Cop 6** |
|---|---|---|---|---|---|
| Latency to 1$^{st}$ paracopulatory | Sub | 67.98 ± 6.26$^{*a}$ | 18.01 ± 6.26$^{b}$ | 14.57 ± 6.26$^{b}$ | 4.34 ± 6.26$^{b}$ |
| | Fully | 20.52 ± 6.26 | 6.07 ± 6.26 | 5.90 ± 6.26 | 2.12 ± 6.26 |
| Mounts | Sub | 0.67 ± 1.17* | 2.75 ± 1.17* | 1.92 ± 1.17 | 2.75 ± 1.17* |
| | Fully | 5.42 ± 1.17$^{ab}$ | 6.08 ± 1.17$^{ab}$ | 3.50 ± 1.17$^{a}$ | 7.92 ± 1.17$^{b}$ |
| Intromissions | Sub | 0.25 ± 1.85$^{*a}$ | 8.75 ± 1.85$^{*b}$ | 6.00 ± 1.85$^{*b}$ | 8.42 ± 1.85$^{*b}$ |
| | Fully | 9.33 ± 1.85$^{a}$ | 16.92 ± 1.85$^{b}$ | 12.83 ± 1.85$^{ab}$ | 16.08 ± 1.85$^{b}$ |
| Ejaculation | Sub | -1.78E+-15±0.35$^{a}$ | 1.33 ± 0.35$^{b}$ | 1.000 ± 0.35$^{*ab}$ | 1.42 ± 0.35$^{b}$ |
| | Fully | 0.58 ± 0.35$^{a}$ | 2.17 ± 0.35$^{b}$ | 2.50 ± 0.53$^{b}$ | 2.25 ± 0.35$^{b}$ |
| Latency to 1st ejaculation | Sub | 1800 ± 162.90$^{*a}$ | 1023.90 ± 162.90$^{*b}$ | 1232.65 ± 162.90$^{*b}$ | 1093.53 ± 162.90$^{*b}$ |
| | Fully | 1326.78 ± 162.90$^{a}$ | 495.23 ± 162.90$^{b}$ | 380.55 ± 162.90$^{b}$ | 506.00 ± 162.90$^{b}$ |
| Percentage of exits after mounts | Sub | 18.57 ± 15.19 | 36.86 ± 10.04 | 24.14 ± 10.49 | 22.33 ± 10.04 |
| | Fully | 1.59 ± 9.47 | 15.61 ± 8.95 | 18.06 ± 8.95 | 11.67 ± 8.95 |
| Percentage of exits after intromissions | Sub | 12.94 ± 25.98$^{ab}$ | 17.26 ± 9.93$^{a}$ | 60.48 ± 9.96$^{b}$ | 61.23 ± 9.62$^{b}$ |
| | Fully | 2.07 ± 10.20$^{a}$ | 24.51 ± 9.32$^{c}$ | 49.31 ± 9.32$^{bd}$ | 44.22 ± 9.32$^{b}$ |
| Percentage of exits after ejaculations | Sub | - | 47.13 ± 17.11 | 66.67 ± 18.48 | 83.59 ± 17.11 |
| | Fully | 33.36 ± 18.48 | 26.39 ± 13.08 | 48.61 ± 13.07 | 53.47 ± 13.07 |
| Contact return latency after mounts | Sub | 8.93 ± 92.59 | 27.39 ± 46.93 | 114.05 ± 54.12 | 83.14 ± 53.81 |
| | Fully | 33.56 ± 64.77 | 113.93 ± 41.81 | 81.97 ± 46.63 | 85.72 ± 46.56 |
| Contact return latency after intromissions | Sub | - | 56.74 ± 21.64 | 106.62 ± 16.79 | 79.35 ± 17.69 |
| | Fully | 70.44 ± 37.34 | 86.76 ± 21.64 | 108.27 ± 17.69 | 56.12 ± 16.79 |
| Contact return latency after ejaculations | Sub | - | 66.33 ± 54.28$^{*a}$ | 323.65 ± 54.28$^{b}$ | 173.97 ± 48.57$^{ab}$ |
| | Fully | 111.63 ± 76.74 | 234.99 ± 54.26 | 269.98 ± 41.04 | 209.94 ± 41.04 |
| Paracopulatory per sexual bout | Sub | 2.66 ± 0.33* | 3.15 ± 0.34* | 2.75 ± 0.34* | 3.29 ± 0.33* |
| | Fully | 5.29 ± 0.33$^{a}$ | 4.70 ± 0.33$^{ab}$ | 3.92 ± 0.33$^{b}$ | 5.22 ± 0.33$^{ac}$ |
| Lordoses per sexual bout | Sub | 0.03 ± 0.05$^{*a}$ | 0.20 ± 0.05$^{ab}$ | 0.25 ± 0.05$^{b}$ | 0.22 ± 0.05$^{*b}$ |
| | Fully | 0.25 ± 0.05 | 0.32 ± 0.05 | 0.28 ± 0.05 | 0.40 ± 0.05 |
| Copulations per sexual bout | Sub | 0.03 ± 0.05$^{*a}$ | 0.22 ± 0.06$^{ab}$ | 0.27 ± 0.06$^{b}$ | 0.25 ± 0.05$^{*b}$ |
| | Fully | 0.26 ± 0.05 | 0.35 ± 0.05 | 0.31 ± 0.05 | 0.44 ± 0.05 |

Note: Data shown as mean ± SEM. *p ≤ 0.05 different with fully-primed females rats of the same Cop test. $^{ab}$ values within a row with unlike letters significantly differed between Cop tests. $p < 0.05$. Cop = copulation test.

**Supplementary table 4.** Results of the statistical mixed models test in Experiment part 2.

| Parameter | N | Statistics | F value | Post Hoc comparison |
|---|---|---|---|---|
| Time spent in the male compartment | Sub-primed N = 10, fully-primed N = 12 | Mixed model | Treatment: F(1, 20) = 0.408, p = 0.530 Drug: F(1, 20) = 0.012, p = 0.915 Treatment x Drug: F(1, 20) = 0.000, p = 0.995 | Effect of drug FP: VEH vs DPAT, p = 0.941 SP: VEH vs DPAT, p = 0.939 Effect of treatment DPAT: FP vs SP, p = 0.659 VEH: FP vs SP, p = 0.665 |
| Total number of paracopulatory behaviors | Sub-primed N = 10, fully-primed N = 12 | Mixed model | Treatment: F(1, 20) = 9.881, p = 0.005 Drug: F(1, 20) = 75.483, p < 0.001 Treatment x Drug: F(1, 20) = 8.565, p = 0.008 | Effect of drug FP: VEH vs DPAT, p < 0.001 SP: VEH vs DPAT, p < 0.001 Effect of treatment DPAT: FP vs SP, p = 0.752 VEH: FP vs SP, p < 0.001 |
| Lordosis quotient | Sub-primed N = 10, fully-primed N = 12 | Mixed model | Treatment: F(1, 14.086) = 0.627, p = 0.441 Drug: F(1, 13.873) = 102,356, p < 0.001 Treatment x Drug: F(1, 13.873) = 0.846, p = 0.373 | Effect of drug FP: VEH vs DPAT, P < 0.001 SP: VEH vs DPAT, P < 0.001 Effect of treatment DPAT: FP vs SP, p = 0.237 VEH: FP vs SP, P = 0.990 |
| Lordosis score | Sub-primed N = 10, fully-primed N = 12 | Mixed model | Treatment: F(1, 20.816) = 0.001, p = 0.977 Drug: F(1, 17.813) = 74.978, p < 0.001 Treatment x Drug: F(1, 17.813) = 5.383, p = 0.032 | Effect of drug FP: VEH vs DPAT, P < 0.001 SP: VEH vs DPAT, P < 0.001 Effect of treatment DPAT: FP vs SP, P = 0.191 VEH: FP vs SP, P = 0.113 |
| Total number of received copulations | Sub-primed N = 10, fully-primed N = 12 | Mixed model | Treatment: F(1, 20) = 15.700, p < 0.001 Drug: F(1, 20) = 18.288, p < 0.001 Treatment x Drug: F(1, 20) = 1.183, p = 0.290 | Effect of drug FP: VEH vs DPAT, P < 0.001 SP: VEH vs DPAT, P < 0.043 Effect of treatment DPAT: FP vs SP, P = 0.008 VEH: FP vs SP, P < 0.001 |
| Mounts | Sub-primed N = 10, fully-primed N = 12 | Mixed model | Treatment: F(1, 20.000) = 4.853, p = 0.039 Drug: F(1, 20.00) = 9.198, p = 0.007 Treatment x Drug: F(1, 20.00) = 0.611, p = 0.444 | Effect of drug FP: VEH vs DPAT, P = 0.010 SP: VEH vs DPAT, P = 0.143 Effect of treatment DPAT: FP vs SP, P = 0.031 VEH: FP vs SP, P = 0.187 |
| Intromissions | Sub-primed N = 10, fully-primed N = 12 | Mixed model | Treatment: F(1, 20.000) = 25.371, p < 0.001 Drug: F(1, 20.00) = 61.411, p < 0.001 Treatment x Drug: F(1, 20.00) = 4.358, p = 0.050 | Effect of drug FP: VEH vs DPAT, P < 0.001 SP: VEH vs DPAT, P < 0.001 Effect of treatment DPAT: FP vs SP, P = 0.041 VEH: FP vs SP, P < 0.001 |
| Ejaculations | Sub-primed N = 10, fully-primed N = 12 | Mixed model | Treatment: F(1, 20.000) = 5.073, p = 0.036 Drug: F(1, 20.00) = 25.794, p < 0.001 Treatment x Drug: F(1, 20.00) = 0.993 , p = 0.331 | Effect of drug FP: VEH vs DPAT, P < 0.001 SP: VEH vs DPAT, P = 0.012 Effect of treatment DPAT: FP vs SP, P = 0.405 VEH: FP vs SP, P = 0.028 |

| | | | | |
|---|---|---|---|---|
| Percentage of exits after mounts | Sub-primed N = 10, fully-primed N = 12 | Mixed model | Treatment:<br>F(1, 15.976) = 9.564, p = 0.007<br>Drug:<br>F(1, 16.064) = 8.066, p = 0.012<br>Treatment x Drug:<br>F(1, 16.064) = 1.020, p = 0.328 | Effect of drug<br>FP: VEH vs DPAT, P = 0.181<br>SP: VEH vs DPAT, P = 0.021<br>Effect of treatment<br>DPAT: FP vs SP, P = 0.005<br>VEH: FP vs SP, P = 0.168 |
| Percentage of exits after intromissions | Sub-primed N = 10, fully-primed N = 12 | Mixed model | Treatment:<br>F(1, 18.870) = 1.009, p = 0.328<br>Drug:<br>F(1, 19.304) = 0.647, p = 0.431<br>Treatment x Drug:<br>F(1, 19.304) = 0.002, p = 0.961 | Effect of drug<br>FP: VEH vs DPAT, P = 0.546<br>SP: VEH vs DPAT, P = 0.595<br>Effect of treatment<br>DPAT: FP vs SP, P = 0.551<br>VEH: FP vs SP, P = 0.421 |
| Percentage of exits after ejaculations | Sub-primed N = 10, fully-primed N = 12 | Mixed model | Treatment:<br>F(1, 4.462) = 0.020, p = 0.894<br>Drug:<br>F(1, 4.462) = 1.036, p = 0.361<br>Treatment x Drug:<br>F(1, 4462) = 0.020, p = 0.894 | Effect of drug<br>FP: VEH vs DPAT, P = 0.333<br>SP: VEH vs DPAT, P = 0.642<br>Effect of treatment<br>DPAT: FP vs SP, P = 1.000<br>VEH: FP vs SP, P = 0.739 |
| Contact return latency after mounts | Sub-primed N = 10, fully-primed N = 12 | Mixed model | Treatment:<br>F(1, 16.260) = 0.282, p = 0.602<br>Drug:<br>F(1, 5.188) = 9.306, p = 0.027<br>Treatment x Drug:<br>F(1, 5.188) = 9.306, p = 0.027 | Effect of drug<br>FP: VEH vs DPAT, P = 0.008<br>SP: VEH vs DPAT, P = 0.999<br>Effect of treatment<br>DPAT: FP vs SP, P = 0.105<br>VEH: FP vs SP, P = 0.570 |
| Contact return latency after intromissions | Sub-primed N = 10, fully-primed N = 12 | Mixed model | Treatment:<br>F(1, 13.090) = 0.354, p = 0.562<br>Drug:<br>F(1, 12.214) = 2.143, p = 0.168<br>Treatment x Drug:<br>F(1, 12.214) = 1.927, p = 0.190 | Effect of drug<br>FP: VEH vs DPAT, P = 0.953<br>SP: VEH vs DPAT, P = 0.091<br>Effect of treatment<br>DPAT: FP vs SP, P = 0.251<br>VEH: FP vs SP, P = 0.485 |
| Contact return latency after ejaculations | Sub-primed N = 10, fully-primed N = 12 | Mixed model | Treatment:<br>F(1, 3.538) = 0.392, p = 0.569<br>Drug:<br>F(1, 16.165) = 0.194, p = 0.665<br>Treatment x Drug:<br>F(1, 16.165) = 0.158, p = 0.696 | Effect of drug<br>FP: VEH vs DPAT, P = 0.379<br>SP: VEH vs DPAT, P = 0.981<br>Effect of treatment<br>DPAT: FP vs SP, P = 0.932<br>VEH: FP vs SP, P = 0.288 |
| Total number of sexual bouts | Sub-primed N = 10, fully-primed N = 12 | Mixed model | Treatment:<br>F(1, 20) = 9.485, p = 0.006<br>Drug:<br>F(1, 20) = 124.269, p < 0.001<br>Treatment x Drug:<br>F(1, 20) = 4.338, p = 0.050 | Effect of drug<br>FP: VEH vs DPAT, P < 0.001<br>SP: VEH vs DPAT, P < 0.001<br>Effect of treatment<br>DPAT: FP vs SP, P = 0.4860<br>VEH: FP vs SP, P < 0.001 |
| Mean duration of sexual bouts | Sub-primed N = 10, fully-primed N = 12 | Mixed model | Treatment:<br>F(1, 20) = 0.267, p = 0.611<br>Drug:<br>F(1, 20) = 9.615, p = 0.006<br>Treatment x Drug:<br>F(1, 20) = 6.132, p = 0.022 | Effect of drug<br>FP: VEH vs DPAT, P < 0.001<br>SP: VEH vs DPAT, P = 0.677<br>Effect of treatment<br>DPAT: FP vs SP, P = 0.178<br>VEH: FP vs SP, P = 0.041 |
| Mean duration of time-outs | Sub-primed N = 10, fully-primed N = 12 | Mixed model | Treatment:<br>F(1, 20) = 0.433, p = 0.518<br>Drug:<br>F(1, 20) = 10.743, p = 0.004<br>Treatment x Drug:<br>F(1, 20) = 1.157, p = 0.295 | Effect of drug<br>FP: VEH vs DPAT, P = 0.004<br>SP: VEH vs DPAT, P = 0.152<br>Effect of treatment<br>DPAT: FP vs SP, P = 0.227<br>VEH: FP vs SP, P = 0.767 |

**Supplementary table 5.** The effect of (R)-(+)-8-OH-DPAT on female sexual behavior.

| **Parameters** | **Sub-primed female rats** | | **Fully-primed female rats** | |
|---|---|---|---|---|
| | **Vehicle** | **DPAT** | **Vehicle** | **DPAT** |
| Mounts | 2.90 ± 1.70 | 5.90 ± 1.70* | 6.00 ± 1.55^ | 11.08 ± 1.55 |
| Intromissions | 8.00 ± 1.28*^ | 1.00 ± 1.28* | 16.75 ± 1.17^ | 4.67 ± 1.19 |
| Ejaculations | 1.50 ± 0.35*^ | 0.10 ± 0.35 | 2.58 ± 0.32^ | 0.50 ± 0.32 |
| Percentage of exits after mounts | 18.622 ± 6.60^ | 41.87 ± 6.22* | 6.43 ± 5.63 | 17.48 ± 5.39 |
| Percentage of exits after intromissions | 57.24 ± 11.56 | 46.96 ± 14.62 | 45.05 ± 9.44 | 35.97 ± 10.90 |
| Percentage of exits after ejaculations | 86.11 ± 10.73 | 100.00 ± 26.29 | 81.67 ± 7.59 | 100.00 ± 11.76 |
| Contact return latency after mounts | 97.56 ± 66.37 | 97.48 ± 59.69 | 44.63 ± 63.31^ | 235.88 ± 54.18 |
| Contact return latency after intromissions | 102.46 ± 18.67 | 37.49 ± 30.48 | 84.65 ± 16.70 | 82.93 ± 23.61 |
| Contact return latency after ejaculations | 178.42 ± 39.13 | 176.03 ± 83.50 | 230.27 ± 26.38 | 184.11 ± 37.93 |

Note: *Mean (± SEM ) significantly different between groups (i.e. sub-primed vs fully-primed). ^Mean (± SEM) values within a row were significantly different between treatments (i.e. vehicle vs DPAT). $p \leq 0.05$. DPAT, P = (R)-(+)-8-OH-DPAT

**Supplementary table 6.** Results of the statistical mixed models test in Experiment part 3.

| Parameter | N | Statistics | F value | Post Hoc comparison |
|---|---|---|---|---|
| Total number of paracopulatory behaviors | Sub-primed N = 12, fully-primed N = 12 | Mixed model | Treatment:<br>F(1, 22) = 25.095, p < 0.001<br>Pacing:<br>F(1, 22) = 10.245, p = 0.004<br>Treatment x Pacing:<br>F(1, 22) = 0.328, p = 0.573 | Effect of setup<br>FP: NPM vs PM, p = 0.014<br>SP: NPM vs PM, p = 0.077<br>Effect of treatment<br>NPM: FP vs SP, P < 0.001<br>PM: FP vs SP, p = 0.002 |
| Lordosis quotient | Sub-primed N = 12, fully-primed N = 12 | Mixed model | Treatment:<br>F(1, 18.758) = 1.735, p = 0.204<br>Pacing:<br>F(1, 18.647) = 0.329, p = 0.573<br>Treatment x Pacing:<br>F(1, 18.647) = 2.145, p = 0.160 | Effect of setup<br>FP: NPM vs PM, p = 0.532<br>SP: NPM vs PM, p = 0.171<br>Effect of treatment<br>NPM: FP vs SP, P = 0.999<br>PM: FP vs SP, p = 0.059 |
| Lordosis score | Sub-primed N = 12, fully-primed N = 12 | Mixed model | Treatment:<br>F(1, 22.311) = 12.254, p = 0.002<br>Pacing:<br>F(1, 22.021) = 10.541, p = 0.004<br>Treatment x Pacing:<br>F(1, 22.021) = 3.455, P = 0.076 | Effect of setup<br>FP: NPM vs PM, p = 0.331<br>SP: NPM vs PM, p = 0.002<br>Effect of treatment<br>NPM: FP vs SP, P < 0.001<br>PM: FP vs SP, p = 0.106 |
| Total number of received copulations | Sub-primed N = 12, fully-primed N = 12 | Mixed model | Treatment:<br>F(1, 22) = 8.139, p = 0.009<br>Pacing:<br>F(1, 22) = 12.098, p = 0.002<br>Treatment x Pacing:<br>F(1, 22) = 0.387, p = 0.540 | Effect of setup<br>FP: NPM vs PM, p = 0.056<br>SP: NPM vs PM, p = 0.008<br>Effect of treatment<br>NPM: FP vs SP, p = 0.082<br>PM: FP vs SP, p = 0.013 |
| Total number of sexual bouts | Sub-primed N = 12, fully-primed N = 12 | Mixed model | Treatment:<br>F(1, 22) = 0.697, p = 0.413<br>Pacing:<br>F(1, 22) = 14.678, p < 0.001<br>Treatment x Pacing:<br>F(1, 22) = 8.778, p = 0.007 | Effect of setup<br>FP: NPM vs PM, p = 0.545<br>SP: NPM vs PM, p < 0.001<br>Effect of treatment<br>NPM: FP vs SP, P = 0.404<br>PM: FP vs SP, P = 0.029 |
| Mean duration of sexual bouts | Sub-primed N = 12, fully-primed N = 12 | Mixed model | Treatment:<br>F(1, 22) = 20.462, p < 0.001<br>Pacing:<br>F(1, 22) = 0.730, p = 0.402<br>Treatment x Pacing:<br>F(1, 22) = 3.105, p = 0.092 | Effect of setup<br>FP: NPM vs PM, p = 0.078<br>SP: NPM vs PM, p = 0.527<br>Effect of treatment<br>NPM: FP vs SP, p < 0.001<br>PM: FP vs SP, p = 0.063 |
| Mean duration of time-outs | Sub-primed N = 12, fully-primed N = 12 | Mixed model | Treatment:<br>F(1, 22) = 5.938, p = 0.023<br>Pacing:<br>F(1, 22) = 14.140, p = 0.001<br>Treatment x Pacing:<br>F(1, 22) = 7.927, p = 0.010 | Effect of setup<br>FP: NPM vs PM, p = 0.511<br>SP: NPM vs PM, p < 0.001<br>Effect of treatment<br>NPM: FP vs SP, P = 0.927<br>PM: FP vs SP, p < 0.001 |
| Latency to 1$^{st}$ paracopulatory | Sub-primed N = 12, fully-primed N = 12 | Mixed model | Treatment:<br>F(1, 22) = 3.425, p = 0.078<br>Pacing:<br>F(1, 22) = 1.401, p = 0.249<br>Treatment x Pacing:<br>F(1, 22) = 0.610, p = 0.443 | Effect of setup<br>FP: NPM vs PM, p = 0.778<br>SP: NPM vs PM, p = 0.179<br>Effect of treatment<br>NPM: FP vs SP, p = 0.067<br>PM: FP vs SP, p = 0.428 |

**Supplementary table 7.** The effect of testing conditions on female sexual behavior.

| Parameters | Sub-primed female rats | | Fully-primed female rats | |
|---|---|---|---|---|
| | **PM** | **NPM** | **PM** | **NPM** |
| **Latency to 1st paracopulatory** | 4.34 ± 1.97 | 8.11 ± 1.97 | 2.12 ± 1.97 | 2.89 ± 1.97 |

Note: *Mean (± SEM) significantly different between groups (i.e. sub-primed vs fully-primed). ^Mean (± SEM) values within a row were significantly different between treatments (i.e. PM vs NPM). $p \leq 0.05$. PM, P = paced mating, NPM, P = non-paced mating.
